# Parkin-dependent ubiquitination of TAX1BP1 affects the degradation pathway of defective mitochondria

**DOI:** 10.1101/2025.05.16.654474

**Authors:** Anna Lechado-Terradas, Bianca Lemke, Katharina I. Zittlau, Boris Macek, Philipp J. Kahle

**Affiliations:** Laboratory of Functional Neurogenetics, Department of Neurodegeneration, Hertie Institute for Clinical Brain Research and German Center for Neurodegenerative Diseases, Faculty of Medicine, University of Tübingen, Tübingen, Germany; Quantitative Proteomics Group, Department of Biology, Interfaculty Institute of Cell Biology, University of Tübingen, Tübingen, Germany; Department of Biochemistry, Faculty of Science, University of Tübingen, Tübingen, Germany

**Author notes:** Laboratory for Structural Mass Spectrometry, Department of Biomolecular Sciences, Weizmann Institute of Science, Rehovot, Israel (Katharina I. Zittlau).

**Keywords:** Autophagosome, endosome, mitophagy, parkin, TAX1BP1, ubiquitination

## Abstract

In stressed cells, the recessive Parkinson disease (PD) associated gene products PINK1 and parkin mediate the autophagic removal of damaged mitochondria (mitophagy). Upon mitochondrial membrane potential disruption, PINK1 phosphorylation activates the ubiquitin ligase parkin which ubiquitinates various mitochondrial protein substrates. These feed-forward modifications on the mitochondria surface attract ubiquitin-binding autophagy receptors that target ubiquitinated mitochondria for degradation. Investigating post-translational protein modifications during this process, we detected transient ubiquitination of K549 within the third coiled-coil domain (CC3) of TAX1BP1 in HeLa cells expressing wild-type (WT) but not catalytically inactive parkin. Parkin-dependent ubiquitination did not target TAX1BP1 to proteasomal degradation but was rather indicative of a regulatory modification. In cells with the full complement of autophagy receptors, TAX1BP1 plays only a minor role in mitophagy. However, when expressed as a sole autophagy receptor, both WT and ubiquitination deficient TAX1BP1 were capable of promoting mitophagy, albeit mitochondria degradation was slightly delayed under mutant conditions. Use of the lysosomal inhibitor bafilomycin A indicated classical autophagolysosomal targeting of damaged mitochondria mediated by WT TAX1BP1. However, for the ubiquitination-deficient TAX1BP1, we observed an increased prevalence of enlarged endolysosomal vesicles carrying accumulated TAX1BP1-positive autophagosomes filled with mitochondrial material. Thus, while ubiquitination of the CC3 domain of TAX1BP1 is not essential for complete mitophagy, the lack of CC3 in TAX1BP1 reroutes the degradation flux to a less efficient endolysosmal degradative pathway. Interestingly, the PD gene product VPS35 becomes prominently engaged in this alternative mitophagy pathway.

## Introduction

Mitochondrial quality control and turnover are essential for cellular homeostasis. Cellular stress associated with mitochondrial membrane depolarization induces a pathway leading to removal of damaged mitochondria activated by the two most common recessive Parkinson disease (PD) associated gene products, PINK1 (phosphatase and tensin homolog induced kinase 1) and PRKN (E3 ligase parkin) [1–4]. Under basal conditions, the nuclear encoded PINK1 protein is efficiently imported into mitochondria, where it is rapidly degraded. Upon dissipation of the mitochondrial membrane potential, PINK1 lacks the driving force for mitochondrial import and hence accumulates on the mitochondrial outer membrane (OMM). There, the PINK1 kinase domain faces the cytosol to phosphorylate ubiquitin and the ubiquitin-like domain of parkin, a process which activates the ubiquitin ligase parkin recruited to the OMM of depolarized mitochondria [5–8]. Various ubiquitin linkages target proteins from depolarized mitochondria to proteasomal and autophagic degradation [9]. Five major selective autophagy receptors have been implicated to connect ubiquitinated mitophagy cargo with the autophagy machinery, including SQSTM1/p62 (sequestosome-1), OPTN (optineurin), CALCOCO2/NDP52 (calcium binding and coiled-coil domain 2), NBR1 (NBR1 autophagy cargo receptor), and TAX1BP1 (Tax1 binding protein 1) [2,10–12]. These selective autophagy receptors do not statically bridge ubiquitinated mitophagy cargo with membranes containing the lipidated form of LC3B (microtubule-associated protein 1 light chain 3 beta), but can be dynamically regulated by post-translational modifications (PTMs). These PTMs are critical modulators of their function and contribute to the recruitment of autophagy-related proteins involved in autophagosome biogenesis (i.e: ATG9 vesicles or the unc-51 like autophagy activating kinase 1 complex) [13–16]. For example, phosphorylation by TANK binding kinase 1 contributes to important regulatory dynamics of OPTN and NDP52 for selective autophagic removal of damaged mitochondria [10,12,16].

Investigating the phospho-proteome and ubiquitylome throughout the time course of mitophagy in catalytically active parkin expressing HeLa cells [17], we detected transient ubiquitination of TAX1BP1 at K549 within the third coiled-coil domain (CC3) of this autophagy receptor. Validation of TAX1BP1 ubiquitination in cells expressing wild-type (WT) or a ligase-dead parkin mutant (C431A), confirmed parkin-dependent ubiquitination of the autophagy receptor at early stages of mitophagy. Removal of the highly ubiquitinated CC3 domain of TAX1BP1 reduced TAX1BP1 distribution on mitochondrial material and induced the engulfment of accumulated autophagosomes into enlarged endolysosomal vesicles (potential amphisomes), apparently rerouting the mitophagy pathway to the endolysosomal system. Thus, the ubiquitinatable CC3 of TAX1BP1 directs depolarized mitochondrial material to autophagosomal degradation, whereas CC3 perturbation mediates the degradation of accumulated autophagosomes with damaged mitochondrial cargo through a less efficient pathway involving endolysosomal trafficking into enlarged endolysosomes. Interestingly, the PD gene product VPS35 (VPS35 retromer complex component) became prominently engaged in the alternative mitophagy pathway described in this study, clearly recognizing the surroundings of the identified enlarged degradative vesicles.

## Results

### Transient ubiquitination of TAX1BP1 in Parkin-dependent mitophagy

Examining the ubiquitylome of HeLa cells stably expressing parkin treated with the mitochondrial uncoupler carbonyl cyanide m-chlorophenylhydrazone (CCCP) [17], we detected transient ubiquitination of TAX1BP1 at K549 (**Figure 1A and B**). The ubiquitination-indicating di-glycine motifs were detected by mass spectrometry predominantly for TAX1BP1 peptides containing K549, with only minor signals for K571, also located in the CC3 region, and K439, upstream of CC3 (**Figure 1A and B**). TAX1BP1 ubiquitination only occurred in cells expressing WT parkin, but not the catalytically inactive form (C431A-parkin) (**Figure 1B**). Other studies previously reported K549-TAX1BP1 to be increasingly ubiquitinated in a parkin-dependent manner and to follow a similar time course as the one described here [18]. Validation of parkin-dependent ubiquitination of TAX1BP1 via western blotting confirmed the differences between WT- and C431A-parkin observed in the mass-spectrometry screen (**Figure 1C**). In HeLa cells stably expressing active parkin, but not ligase-inactive C431A-parkin, CCCP treatment caused a band shift compatible with TAX1BP1 poly-ubiquitination peaking around 4 h after mitochondrial membrane depolarization. TAX1BP1 ubiquitination vanished at later time points of mitophagy, when parkin substrates like TOM20 (translocase of outer mitochondrial membrane 20) and VDAC (voltage dependent anion channel) were degraded (**Figure 1C**).

**Figure 1:**
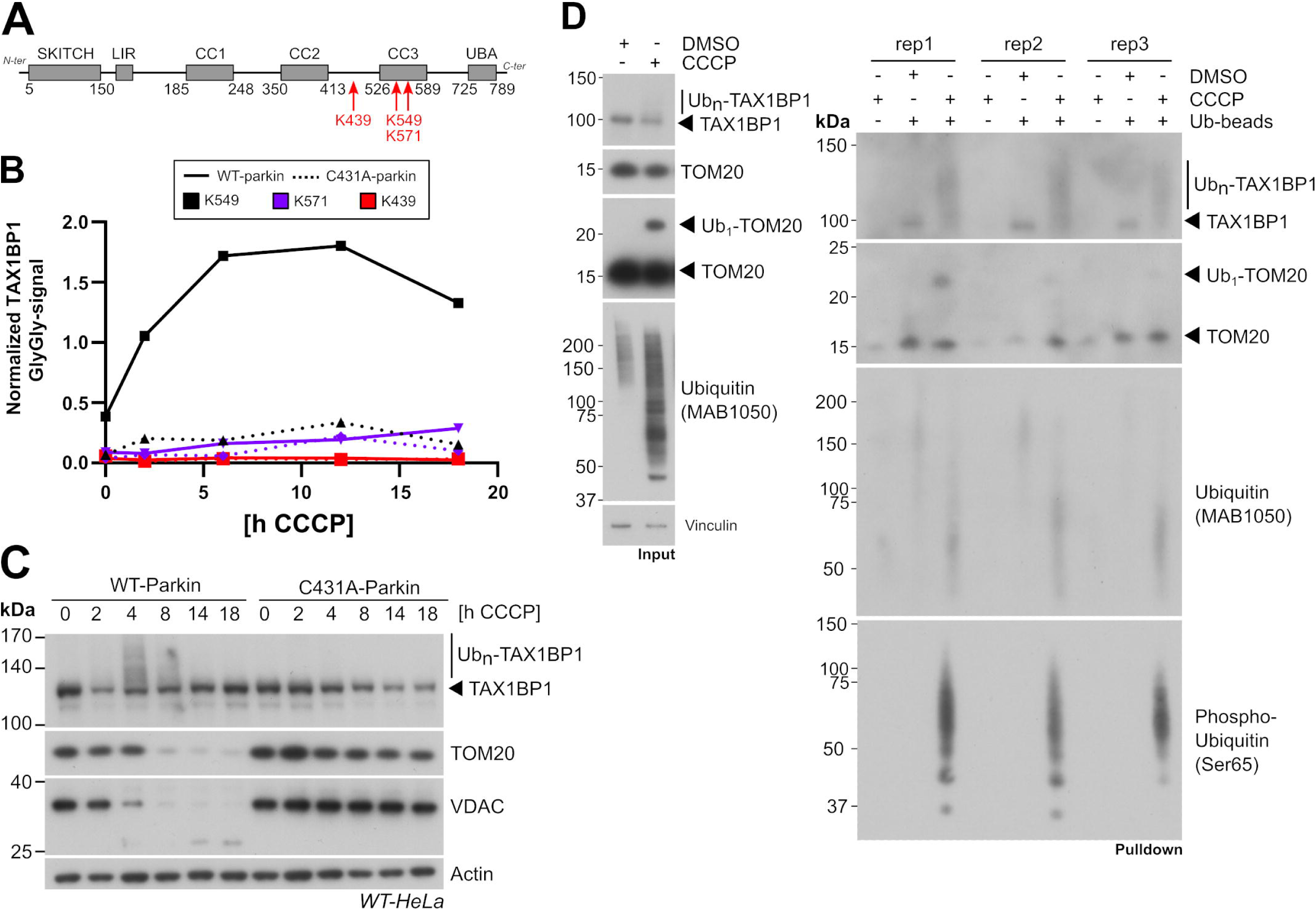
Ubiquitination of TAX1BP1 upon CCCP-induced mitophagy in HeLa cells stably expressing WT- or C431A-Parkin. (**A**) Schematic representation of TAX1BP1 domain structure. Identified ubiquitinated lysine residues from the ubiquitylome mass-spectrometry screen are indicated in red. (**B**) Normalized Gly-Gly peptide signals of the identified ubiquitinated residues in both WT- and C431A-parkin HeLa cell lines during mitophagy induction at the indicated time points. (**C**) Western blot detection of the indicated proteins in whole cell lysates from HeLa cells stably expressing WT- or catalytically inactive C431A-parkin treated with CCCP for the indicated times. TAX1BP1 band shift peaking at 4 h confirms ubiquitination. (**D**) Western blots showing protein signals of 5% of input or 10% of eluted pulldown samples performed in WT-Parkin Hela depolarized with CCCP for 4 h or under control conditions (DMSO). Pulldown samples are shown for three independent replicate experiments (Rep).

To confirm that the observed TAX1BP1 band shifts reflect ubiquitination of the autophagy adaptor, ubiquitin pulldown experiments were performed in the WT-Parkin HeLa cell line (**Figure 1D**). After short CCCP depolarization, some parkin-dependent substrates were pulled down such as ubiquitinated (Ub-) TOM20 as well as ubiquitin in its unmodified form (Total Ub) or the Ser65-phosphorylated version (pS65-Ub) (**Figure 1D**). The appearance of a smear-like band shift for TAX1BP1 was also detected under the same conditions, which contrasted with the absence of a smear observed under basal or bead control pulldown conditions (**Figure 1D**). This data confirms the ubiquitination nature of endogenousTAX1BP1 occurring in a parkin-dependent mitophagy context.

Overall, TAX1BP1 levels decreased parallelly to VDAC and TOM20 degradation during early depolarization (2-4 h CCCP) but increased after extended depolarization (8-18 h CCCP) (**Figure 1C**), indicating that early steps of mitophagy might also involve some degradation of the autophagy receptor along with mitochondrial cargo. As early stages of PINK1/parkin mitophagy rely on proteasomal degradation of immediate-early parkin substrates, we checked if the identified ubiquitination of TAX1BP1 could target the autophagy receptor for early proteasomal degradation. In order to test this, HeLa cells stably expressing parkin were transfected with TAX1BP1 and subjected to mitophagy in the presence or absence of the proteasomal inhibitor MG132 (**Fig. S1**). Here, while endogenous TAX1BP1 was not reduced after short depolarization (2 h CCCP), a slight reduction of overexpressed TAX1BP1 levels was observed at this early time point (**Fig. S1**). Importantly, proteasome inhibition did not stabilize poly-ubiquitinated TAX1BP1, neither in the endogenous or overexpressed form, contrary to what was observed for the known proteasome target MFN1 (mitofusin 1) (**Fig. S1**). Thus, the rapid TAX1BP1 ubiquitination observed in this study, seemed to not be part of the initial proteasome-targeting ubiquitination program of OMM-associated proteins, but rather appears to be a regulatory PTM of TAX1BP1.

### Ubiquitinated TAX1BP1 is enriched in depolarized mitochondria

Mitochondria depolarization triggers the activation of parkin-dependent ubiquitination of mitochondrial proteins that will be recognized by the ubiquitin binding domains of autophagy receptors for further successful delivery of mitochondrial cargo to autophagosomes [19]. As expected, endogenous TAX1BP1 translocated to mitochondria after depolarization (**Figure 2A**). To investigate if the ubiquitinated form of TAX1BP1 was also present on mitochondria, sucrose fractionation experiments were performed. After sucrose fractionation, mitochondrial proteins (i.e: TOM20, VDAC and CS [citrate synthase]) were highly enriched in the mitochondria fraction (MTF) (**Figure 2B**). Under no stress conditions, TAX1BP1 was enriched in the membrane fraction (MMF), suggesting that TAX1BP1 could localize directly to membranous compartments under basal conditions. However, upon short mitochondrial depolarization, TAX1BP1 was enriched in the MTF, also in the ubiquitinated form (**Figure 2B**). These data indicated that ubiquitinated TAX1BP1 localized on depolarized mitochondria where it may contribute to parkin-dependent mitophagy.

**Figure 2:**
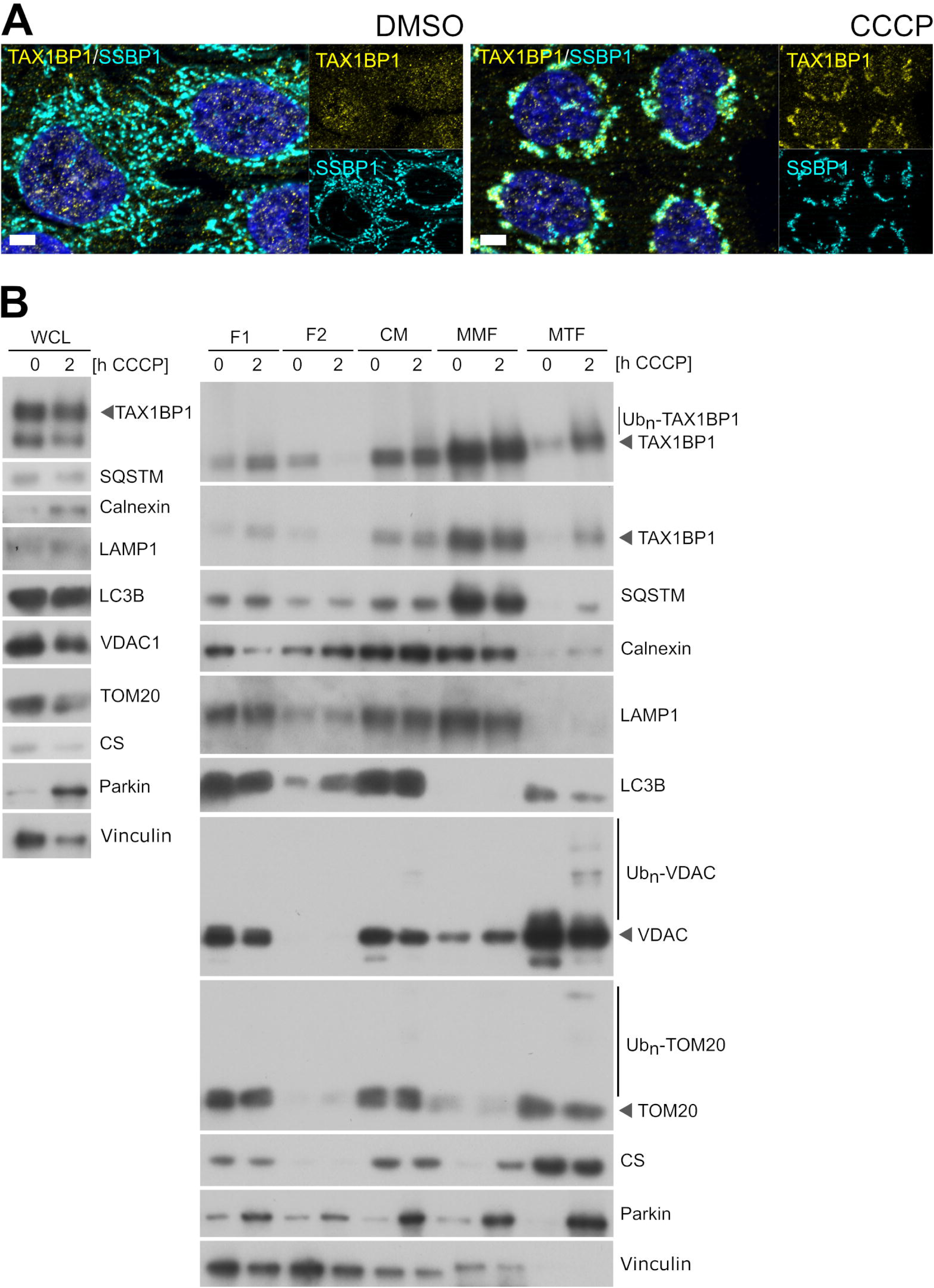
TAX1BP1 translocation to mitochondria during CCCP-induced mitophagy. (**A**) Representative immunofluorescence images showing endogenous TAX1BP1 (yellow), and the inner mitochondrial marker SSBP1 (cyan) at basal levels or after mitophagy induction (10 µM CCCP, 2h). Scale bars: 5 µm (**B**) Subcellular fractionation of WT-Parkin HeLa cells under basal conditions or after short depolarization, where translocation of parkin and ubiquitinated TAX1BP1 to mitochondria can be observed. WCL: whole cell lysate, F1: fraction 1 corresponds to the supernatant recovered after the first centrifugation series, F2: fraction 2 corresponds to the supernatant recovered after the second centrifugation step in which the crude mitochondria pellet were obtained, CM: crude mitochondria fraction, MMF: membrane enriched fraction, MTF: mitochondrial enriched fraction.

### Loss of the TAX1BP1 ubiquitinatable domain reduces clearance of ubiquitinated TOM20

TAX1BP1 is one of the autophagy receptors capable of recognizing ubiquitinated mitochondria and triggering its selective degradation during PINK1/parkin mitophagy [19]. However, among the identified five selective autophagy receptors involved in mitophagy, OPTN, NDP52 and TAX1BP1 are characterized as the only ones capable of inducing mitophagy on their own, with TAX1BP1 being the least efficient of the three [11]. Our results are in accordance with the hypothesis of TAX1BP1 having a minor role amongst other autophagy receptors, as siRNA mediated knock-down of TAX1BP1 in WT-parkin HeLa cells did not abolish the degradation of the outer mitochondrial proteins TOM20 and VDAC (**Fig. S2A**). On the other hand, overexpression of TAX1BP1 had a positive effect on mitophagy, as we observed an increased mitochondrial clustering upon short depolarization (**Fig. S2B**). These results confirmed the involvement of TAX1BP1 in parkin-mediated mitophagy and strengthened the notion of TAX1BP1 having a minor role when all other autophagy receptors are present. Thus, in order to be able to characterize TAX1BP1 functionality isolated in a null autophagy receptor background, HeLa cells lacking all five major autophagy receptors were used (from now on, referred as 5KO HeLa) [11].

To understand the relevance of the identified TAX1BP1 ubiquitination events, lysine substitution (TAX1BP1^K549R^) and deletion (TAX1BP1ΔCC3) mutants were generated (**Figure 3A**). Here, 5KO HeLa cells were transiently co-transfected with WT-parkin combined with the indicated TAX1BP1 mutants and mitophagy was induced via CCCP-mediated depolarization at the indicated time points. Upon co-expression of WT-parkin and WT or TAX1BP1^K549R^ after early mitochondrial depolarization (4 h CCCP), TAX1BP1 ubiquitination was observed, even in the absence of all other four complementary autophagy receptors (**Figure 3B**). In contrast, ubiquitination was reduced upon expression of TAX1BP1ΔCC3 under the same conditions (**Figure 3B-C**). Likewise, TAX1BP1ΔCC3 ubiquitination was significantly reduced in 5KO HeLa stably expressing parkin (from now on 5KO-PRKN HeLa) (**Figure 3D-E**). In ubiquitin pulldown experiments with 5KO-PRKN cells, positive controls such as Ub-TOM20, pS65-Ub and total-Ub were captured upon short CCCP-mediated depolarization but not under basal or control bead conditions (**Figure 3D**). Importantly, when analyzing TAX1BP1 ubiquitination status in pulldown experiments, additional higher molecular bands were observed for both WT and TAX1BP1^K549R^, while only a residual smear after long exposure was observed for TAX1BP1ΔCC3 (**Figure 3D**). Further quantification of these identified higher molecular bands after long TAX1BP1 exposure revealed a significant ubiquitination reduction in TAX1BP1ΔCC3 only (**Figure 3E**). While we cannot discard that residual ubiquitination also occurs in TAX1BP1ΔCC3 -as it is evidenced by the residual smear observed upon depolarization-, a reduced ubiquitination signal is observed in TAX1BP1ΔCC3 when similar TAX1BP1 intensity bands are compared (**Figure 3D**, red vs. blue arrowheads), and this effect is further confirmed upon quantification (**Figure 3E**). The effects observed for TAX1BP1ΔCC3, contrast with the ubiquitination status observed for TAX1BP1^K549R^, which did not show any significant difference when compared to WT (**Figure 3D-E**). Overall, these data indicate that most of the TAX1BP1 ubiquitination events observed upon mitochondria depolarization were localized at the CC3 domain of the autophagy receptor, but were not restricted to K549. It is important to note that the CC3 domain of TAX1BP1 is particularly lysine-rich compared to CC1 or CC2 and that the lysines encoded in the CC3 domain of TAX1BP1 are highly conserved across species (**Fig. S3**). Thus, it is very possible that the sole substitution of a single lysine (i.e: K549R) in this domain is not enough to have an impact on the total TAX1BP1 ubiquitination levels, as many other lysines could be alternatively ubiquitinated in this condition.

**Figure 3:**
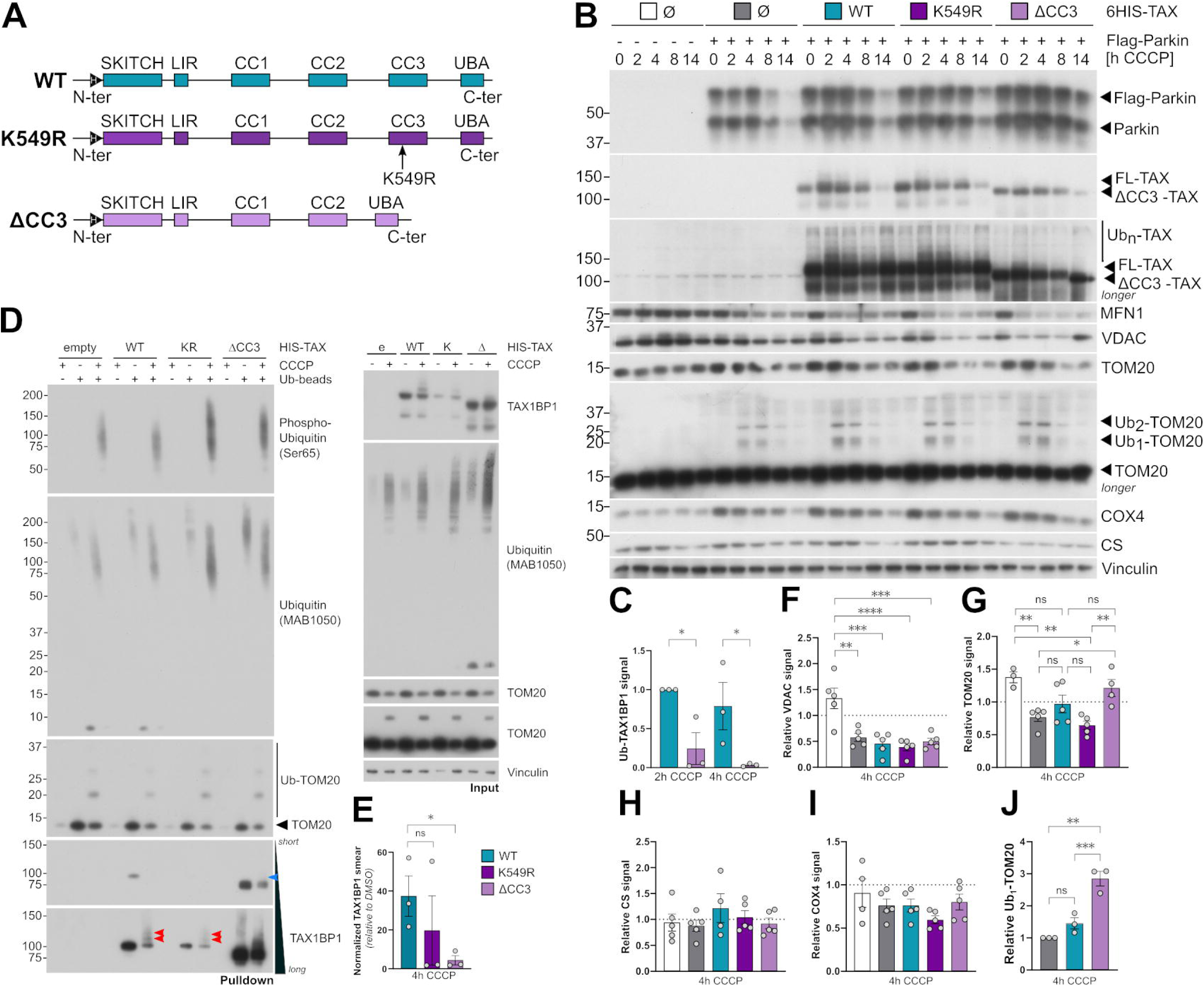
Over-expression of TAX1BP1ΔCC3 leads to changes in its ubiquitination pattern and reduces clearance of ubiquitinated TOM20. (**A**) Schematic representation of the HIS-tagged TAX1BP1 constructs harboring residue-specific or deletion mutations at the CC3 domain. H: 6xHis-tag. (**B**) Mitophagy time course in 5KO-HeLa cells transiently expressing WT, TAX1BP1^K549R^ and TAX1BP1ΔCC3 with or without Flag-tagged WT-Parkin co-transfection. Ø: empty control plasmids. Color-coded legends are shown for each transfection condition. (**C**) Relative levels of ubiquitinated (Ub-) TAX1BP1 after the indicated CCCP treatments. Quantification corresponds to the Ub_n_-TAX1BP1 smear indicated in panel (B). (**D**) Ubiquitin pulldown western blots showing protein signals of 5% of input or 10% of eluted samples performed in 5KO-PRKN HeLa depolarized for 4 h of CCCP compared to control conditions (DMSO). Red and blue arrowheads indicate presence or absence of poly-ubiquitinated TAX1BP1 bands, respectively. (**E**) Quantification of Ub-TAX1BP1 bands obtained in Ubiquitin pulldown shown in (D, lower panel). Ubiquitination levels were normalized to TAX1BP1 input signal for each transfection condition. Further, normalization to control beads and to DMSO control was performed. For this comparison, unpaired t-test was used to assess significance. (**F-I**) Relative protein levels of the following mitochondrial markers: VDAC (F), TOM20 (G), CS (H) and COX4 (I). Quantifications correspond to protein levels in 5KO Hela shown in (B). (**J**) Relative levels of ubiquitinated (Ub-) TOM20 after the indicated CCCP treatment shown in (B). All protein quantifications are derived from at least three independent experiments. Protein signal ratios in (F-I) were obtained after normalization of protein signal with loading control (vinculin) and subsequent normalization with protein values at basal level. Ubiquitination levels shown in panels (C) and (J) were normalized to loading and to TOM20 or TAX1BP1 levels, respectively. Ratios relative to empty transfection control are shown for TOM20 while for Ub-TAX1BP1, ratios are relative to Ub-TAX1BP1-WT levels after 2 h CCCP. One-way ANOVA was used to assess significance unless otherwise specified. Bars: Mean ± SEM.

Next, we focused on investigating if and how the ubiquitination status of TAX1BP1 could affect the degradation of specific mitochondrial proteins. For this purpose, we quantified protein levels of the outer and inner mitochondrial proteins: TOM20, VDAC, CS and COX4 (cytochrome c oxidase subunit 4I1) upon co-expression of WT-Parkin and the different TAX1BP1 mutants after CCCP-mediated depolarization. Under control conditions, where neither TAX1BP1 nor parkin were expressed, protein levels of all analyzed proteins (VDAC, TOM20, CS and COX4) were not significantly reduced after 4 h of CCCP (**Figure 3B and 3F-I**). As expected, upon expression of parkin alone or together with WT TAX1BP, an increase in OMM protein degradation was observed (i.e: TOM20 and VDAC) while inner mitochondrial proteins (COX4 and CS) remained unaffected after 4 h of CCCP, when compared to empty controls (**Figure 3F-I**). These results are in agreement with previous studies indicating that the degradation of inner mitochondrial targets in a parkin-expressing HeLa cell model, require longer depolarization exposure [9,17]. Expression of TAX1BP1^K549R^ did not have any impact on VDAC or TOM20 degradation (**Figure 3F-G**). VDAC degradation was increased upon addition of WT or any TAX1BP1 mutant, when compared to parkin transfection alone. However, changes were found to not be significant, suggesting that the autophagy receptor may not be required for the elimination of this particular OMM protein (**Figure 3B and F**). Similarly, no significant changes in TOM20 degradation were observed upon TAX1BP1 over-expression, as TOM20 degradation was already enhanced with parkin overexpression alone (**Figure 3B and G**), in agreement with previous studies [11]. However, despite no observed changes in protein degradation under these conditions, a significant increase in Ub-TOM20 was identified in TAX1BP1ΔCC3 (**Figure 3G and J**). Thus, these data suggest that the loss of the CC3 ubiquitinatable domain on TAX1BP1 (TAX1BP1ΔCC3) has an impact on the targeting of ubiquitinated TOM20, potentially delaying TOM20 degradation.

### Modulation of the ubiquitination pattern of TAX1BP1 has mild effects on its translocation behavior to mitochondria

The engulfment of mitochondrial material inside autophagosomes is a step that depends on autophagy receptor functionality and that allows: (i) the recognition of ubiquitinated mitochondrial proteins and (ii) the recruitment of autophagy effectors, which will promote autophagosome biogenesis and eventually allow cargo engulfment [20]. Considering this, we hypothesized that the slight defects observed in mitochondrial degradation could be due to a delay in the TAX1BP1-dependent recognition of mitochondrial material. For this purpose, TAX1BP1 translocation behavior and mitochondria clustering were analyzed.

First, basic immunofluorescence was used to visualize TAX1BP1 translocation on mitochondrial material upon co-expression of WT-parkin (**Figure 4A and Fig. S4A**). Under basal conditions, puncta-like structures were observed for all three TAX1BP1 constructs (**Fig. S4A**). Short mitochondria depolarization induced the translocation of all three TAX1BP1 constructs (**Figure 4A**), indicating that both TAX1BP1^K549R^ and TAX1BP1ΔCC3 were capable of recognizing depolarized mitochondria. However, TAX1BP1ΔCC3 translocation to mitochondria was rather localized in specific regions of the SSBP1 cluster and, unlike parkin translocation, did not co-localize with most of SSBP1 material in some cases (**Figure 4A, zoom panel iv**).

**Figure 4:**
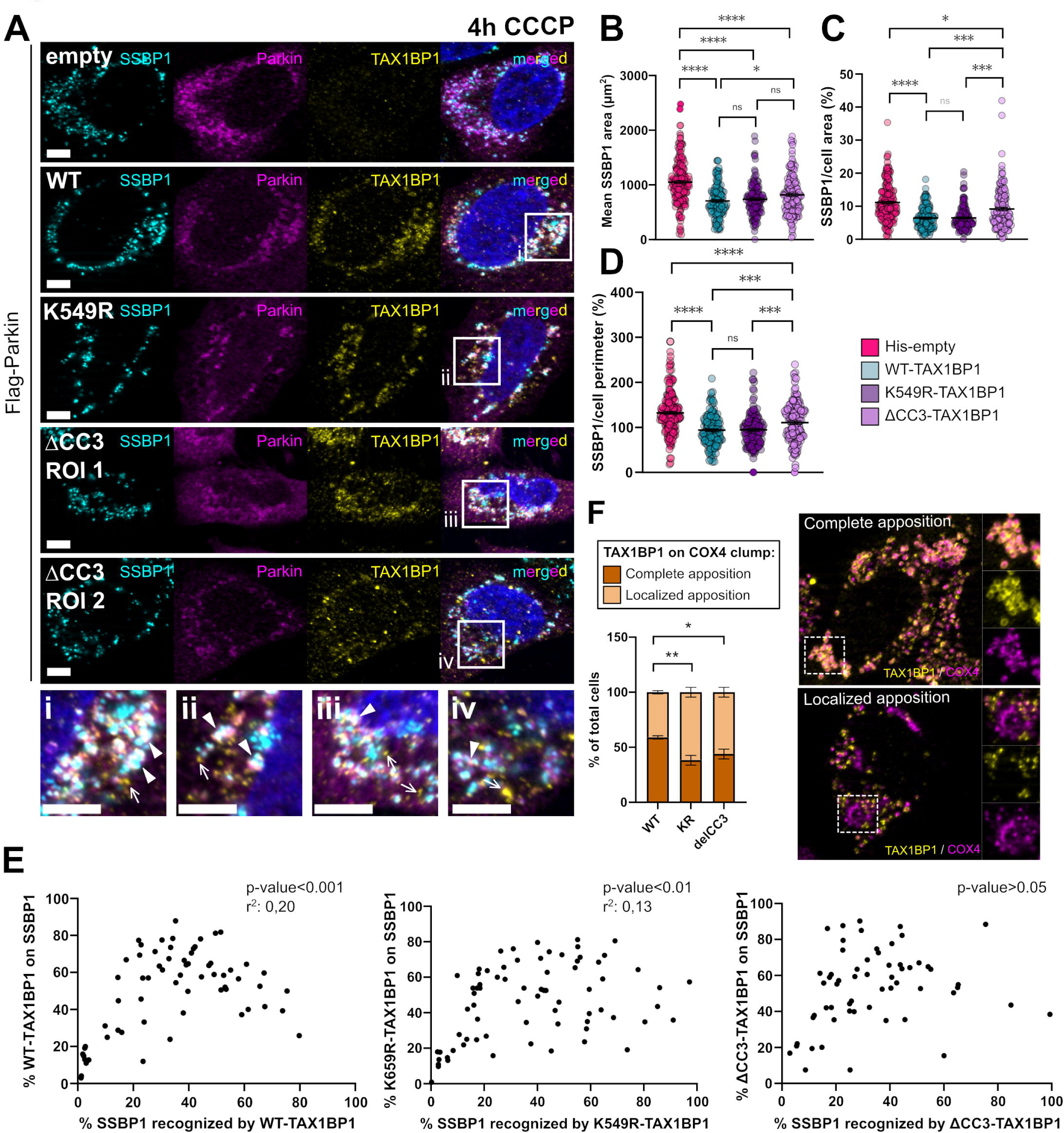
Single cell analysis of TAX1BP1 translocation to mitochondria after CCCP-mediated depolarization. (**A**) Representative immunofluorescence images showing TAX1BP1 (yellow), parkin (magenta) and SSBP1 (cyan) after mitochondria depolarization in 5KO HeLa cells transiently co-expressing Parkin and TAX1BP1. Solid arrowheads indicate TAX1BP1 structures co-localizing with mitochondrial material. Arrows indicate TAX1BP1 structures not engaged with mitochondrial material. ROI 1 represents > 50 % of cell population while ROI 2 represents < 50% of cases. Scale bars: 10 µm. Zoom insets for WT (i), TAX1BP1^K549R^ (ii), TAX1BPΔCC3 (iii and iv) are shown below. (**B**) Mean mitochondrial size of SSBP1 foci after 4h CCCP. (**C-D**) Normalized SSBP1 (C) area or (D) perimeter per cell. (**E**) Correlation analysis of the percentage of mitochondrial area recognized by TAX1BP1 (X axis) in relation to the percentage of total TAX1BP1 found on mitochondria (Y axis). Significance p-values are indicated in the graph for the each TAX1BP1 construct. (**F**) Left panel: quantitative analysis of WT, TAX1BP1^K549R^ or TAX1BP1ΔCC3 localization after mitochondria depolarization in 5KO-PRKN cells after mitophagy induction (4h CCCP). TAX1BP1 localization is classified between cells where COX4 clumps are fully recognized by TAX1BP1 (complete apposition) and cells where COX4 clumps recognition is limited (localized apposition). Bars: mean ± SEM. Right panel: representative immunofluorescence images stained for TAX1BP1 (yellow) and COX4 (magenta) are shown for each category represented in graph.

Next, mitochondrial morphology analysis was performed by quantifying mitochondria area after depolarization. In this case, given that matrix protein degradation was not significantly affected after short depolarization (**Figure 3B and H**), the matrix protein SSBP1 was used for mitochondrial area quantification by using CellProfiler as previously performed [21].

Here, after short depolarization and co-expression of parkin with WT TAX1BP1, we observed a strong reduction of SSBP1 area when compared to control conditions, indicative of mitochondrial clustering and successful mitophagy (**Fig. S4B**). Importantly, while a reduction of mitochondrial area was also observed with parkin expression only, mitochondrial clustering was not as prominent as when WT TAX1BP1 was co-expressed (**Fig. S4B**), indicative of delayed or impaired mitophagy in absence of the autophagy receptor. When analyzing mitochondrial clustering upon co-expression of WT and mutant TAX1BP1, an increase in SSBP1 area and perimeter was observed for TAX1BP1ΔCC3 while no changes were observed for TAX1BP1^K549R^ (**Figure 4B-D**). These data indicated that TAX1BP1ΔCC3 had a delayed efficacy in promoting mitophagy.

Above results seem to indicate that the recognition of depolarized mitochondria is not impaired by any of the mutants, as both TAX1BP1^K549R^ and TAX1BP1ΔCC3 were co-localizing with mitochondrial material upon depolarization (**Figure 4A**). However, upon expression of TAX1BP1ΔCC3 and short depolarization, we observed an increase in Ub-TOM20 (**Figure 3B and J**) as well as a general delay in mitochondrial clustering, indicated by an increase in SSBP1 foci area and perimeter (**Figure 4B-D**). Additionally, a partial TAX1BP1 translocation to specific SSBP1 cluster regions was observed (**Figure 4A, zoom panel iv**). Thus, we hypothesized that the observed TAX1BP1ΔCC3 delay in mitophagy progression may be related to TAX1BP1 capability to recognize SSBP1 clusters. For this purpose, the distribution of TAX1BP1 on depolarized mitochondria material was analyzed with CellProfiler. In general, all TAX1BP1 constructs seemed to recognize SSBP1 material in a similar manner, with distribution percentages ranging from 35-45% of SSBP1 area (**Fig. S4C**). While no significant differences were observed between WT and TAX1BP1 mutants, a slight reduction of the mitochondrial area recognized by TAX1BP1ΔCC3 was observed, when compared to WT and TAX1BP1^K549R^ (**Fig. S4C**). To exclude that the observed differences in TAX1BP1 mitochondrial distribution could be due to distinct mitophagy progression stages, parkin distribution on mitochondrial material was additionally analyzed. In this case, no overall differences were observed upon transfection of WT or TAX1BP1 mutants, as parkin-dependent recognition of SSBP1 area was found between 60-65% for all cases (**Fig. S4D**). We additionally monitored the percentage of total TAX1BP1 involved in the recognition of depolarized mitochondria. Here, no differences were observed for WT and TAX1BP1 mutant, showing 45-50% of total TAX1BP1 to be engaged with depolarized mitochondria (**Fig. S4E**). We further hypothesized that the differences of TAX1BP1 distribution on SSBP1 foci could be caused by differences in TAX1BP1 capability to translocate to mitochondria. For this purpose, the recognized SSBP1 area (X) was plotted against the percentage translocated TAX1BP1 on mitochondria (Y) and correlation analyses were performed for each condition. Indeed, a significant correlation was observed for WT TAX1BP1, with increasing TAX1BP1 being able to recognize 40-50% of SSBP1 area, followed by a TAX1BP1 translocation decrease covering up to 60-80% of SSBP1 area (**Figure 4E**). Correlation analysis was also significant for TAX1BP1^K549R^, which behaved similarly as WT but with an increased variability on the percentage of translocated TAX1BP1^K549R^ (Y axis), after reaching a maximum of 70% translocated TAX1BP1 (**Figure 4E**). Surprisingly, unlike for WT and TAX1BP1^K549R^, no significant correlation was observed for TAX1BP1ΔCC3 (**Figure 4E**). These data suggested an inefficient capability of TAX1BP1ΔCC3 to actively translocate to depolarized mitochondria and thus, distinct binding behavior in comparison to WT TAX1BP1. Additionally, to discard that the differences observed could be due to upstream differences in parkin translocation, a similar correlation analysis was performed for parkin (**Fig. S4F**). Here, significant correlation was found in all cases and no drastic differences were observed for parkin translocation upon expression of all three TAX1BP1 constructs (**Fig. S4F**). In this case, approximately >80% of SSBP1 was found recognized by parkin (**Fig. S4F**), indicating a comparable parkin translocation and thus a similar mitophagy progression stage for all three TAX1BP1 expression cases.

Next, to validate the previously observed differences in TAX1BP1ΔCC3 binding behavior in a context of stable parkin expression, TAX1BP1 mutants were transfected in 5KO-PRKN HeLa cells and TAX1BP1 translocation was analyzed after 4 h of CCCP treatment. To confirm mitophagy progression was at an advanced level, only cells with mitochondrial COX4-positive clusters were taken into account. Here, cells were classified based on TAX1BP1 translocation to mitochondria (complete vs. localized distribution) (**Figure 4F**). Interestingly, a significant increase of cells with localized TAX1BP1 recognition was observed under conditions of TAX1BP1^K549R^ and TAX1BP1ΔCC3 when compared to WT (**Figure 4F**). These data indicated that the recognition of depolarized mitochondria was slightly impaired in TAX1BP1^K549R^ and TAX1BP1ΔCC3 (**Figure 4E and F**).

### Blocking autophagy flux in TAX1BP1ΔCC3 promotes the formation of enlarged TAX1BP1 structures

Next, we sought to examine if the observed differences in mitophagy progression and binding behavior of TAX1BP1^K549R^ and TAX1BP1ΔCC3 would translate in the capability of the autophagy receptor to promote complete elimination of mitochondria. For this purpose, mitochondrial degradation after extended depolarization was analyzed in 5KO-PRKN HeLa cells transfected with WT and TAX1BP1 mutants. Interestingly, all TAX1BP1 constructs were able to induce mitochondria degradation after extended CCCP treatment, as an increased number of cells without mitochondria were observed in comparison to control conditions (**Figure 5A and C**). However, when blocking the autophagy flux with bafilomycin A (BafA), we could observe the presence of small and enlarged TAX1BP1^+^ structures co-localizing with the matrix protein SSBP1 (**Figure 5B**). Importantly, the percentage of cells harboring enlarged TAX1BP1^+^ structures was significantly higher in TAX1BP1ΔCC3 compared to WT or TAX1BP1^K549R^ (**Figure 5D**). In addition, TAX1BP1 ubiquitination status at that time point was validated by means of western blotting. Here, in accordance to what was observed at an earlier stage of mitophagy (**Figure 3B**), ubiquitination of TAX1BP1 was conserved upon autophagy flux blockage for WT and TAX1BP1^K549R^, while a strong ubiquitination reduction was seen for TAX1BP1ΔCC3 under these conditions (**Figure 5E**).

**Figure 5:**
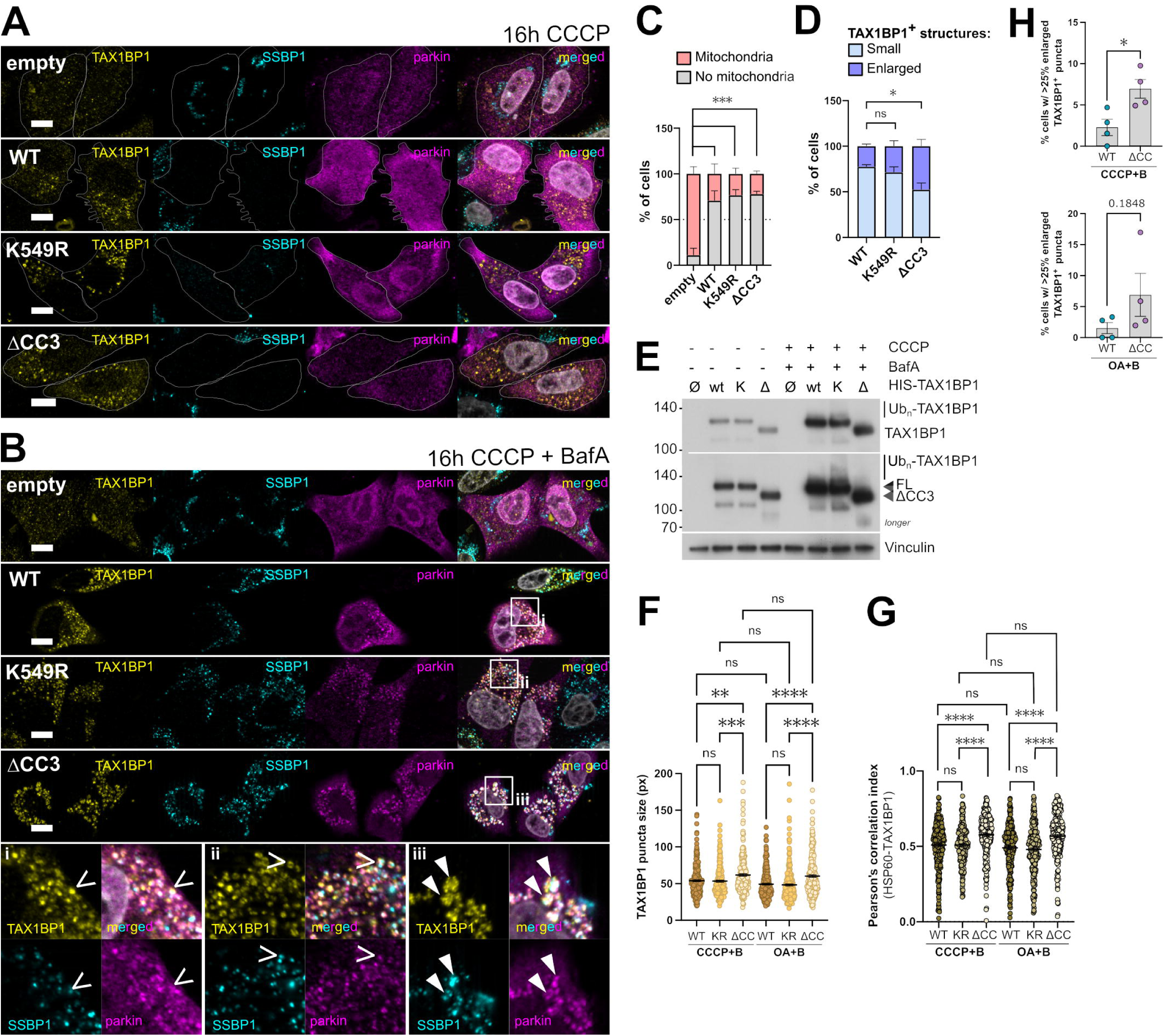
Autophagy flux blockage effects in 5KO-PRKN HeLa cells. (**A-B**) Representative immunofluorescence images of 5KO-PRKN HeLa cells showing TAX1BP1 (yellow), parkin (magenta) and SSBP1 (cyan) after mitochondria depolarization only (A) or with Bafilomycin A (BafA) (B). In (A), the cell contour of cells lacking SSBP1 is highlighted. In (B), zoom insets are shown for WT (i), TAX1BP1^K549R^ (ii) and TAX1BPΔCC3 (iii), where solid arrowheads indicate enlarged TAX1BP1/SSBP1-positive puncta and thin arrowheads indicate small TAX1BP1/SSBP1-positive puncta. (**C-D**) Eye-based quantification of (C) cells with remaining SSBP1 material after extended depolarization (16 h CCCP) and (D) cells harboring enlarged or small TAX1BP1 puncta positive for SSBP1 (16 h CCCP + BafA). (**E**) Changes in TAX1BP1 ubiquitination after 16 h of CCCP treatment combined with BafA. **(F)** Mean TAX1BP1 puncta size after autophagy blockage with BafA combined with CCCP or Oligomycin and Antimycin A (OA) treatment for 16 h. **(G)** Pearson correlation indexes for TAX1BP1 and HSP60 signals after CCCP and OA treatment combined with BafA. In panels (E) and (G), data of individual cells is represented by single dots. All data come from at least three independent experiments. **(H)** Percentages of cells showing at least 25% of TAX1BP1 enlarged puncta after the indicated treatments. Bars: Mean ± SEM. Scale bars: 10 µm. KR: K549R, ΔCC: ΔCC3.

To confirm that the observed phenotype was not derived from CCCP side effects, additional experiments were performed using a combination of Oligomycin and Antimycin A (OA) in addition to the CCCP treatment (**Figure 5F-G and Figure S5).** Here, similarly as observed in previous experiments, we could identify an increase in TAX1BP1ΔCC3 puncta size when compared to WT or TAX1BP1^K549R^ under both treatment conditions (**Figure 5F**). Importantly, the identified TAX1BP1ΔCC3 enlarged puncta were positive for HSP60 (**Figure 5G**). In addition, while TAX1BP1 puncta size significantly increased upon depolarization for both WT and TAX1BP1 mutants, no size differences were observed under BafA control conditions among WT TAX1BP1 and mutants (**Figure S5A-B**).

Next, CellProfiler was used to stratify TAX1BP1 puncta according to their size into enlarged, intermediate or small puncta (**Figure S5C and D**). Such unbiased classification confirms the previous eye-based quantification results (**Figure 5D**). Consistently, an increased intracellular percentage of enlarged TAX1BP1 puncta was observed under TAX1BP1ΔCC3 expression (**Figure S5E**). Following TAX1BP1 puncta size-stratification, cells with more than 25% of enlarged TAX1BP1 puncta were quantified upon CCCP and OA depolarization (**Figure 5H**). Here, a significant increase of cells with enlarged TAX1BP1ΔCC3 puncta was observed under CCCP treatment, while a less prominent increase was identified under OA conditions (**Figure 5H**).

Overall, these data suggested that, while TAX1BP1ΔCC3 was able to promote mitochondria elimination as TAX1BP1^K549R^ and WT TAX1BP1 after long depolarization, the degradative mechanisms driven by TAX1BP1ΔCC3 could differ, as its expression induces a significantly higher prevalence of enlarged TAX1BP1^+^ structures carrying mitochondrial material.

### Enlarged TAX1BP1ΔCC3 structures are not simple autophagosomes

Given the fact that TAX1BP1 is considered the least effective autophagy receptor to be able to initiate mitophagy on its own in a 5KO-HeLa cell model [11], it was tempting to speculate that the enlarged TAX1BP1^+^ structures carrying mitochondrial material could be indicative of mitochondrial degradation occurring via a distinct autophagy pathway. Thus, several markers were used to investigate the nature of these enlarged TAX1BP1^+^ structures. Mitochondrial material was identified by using the matrix markers SSBP1 or HSP60 (60 kDa heat shock protein, mitochondrial). As BafA is known to inhibit autophagic flux, the first marker to test was the autophagosome marker LC3B. The enlarged TAX1BP1^+^ HSP60^+^ structures of WT, TAX1BP1^K549R^ and TAX1BP1ΔCC3 were LC3B^+^ (**Figure 6A**). In all cases, these structures seemed to be of a more complex nature, as they were composed of multiple LC3B^+^ entities structures across the Z plane (**Figure 6B, Fig. S6 and Supplementary videos1-4**). These results suggest the enlarged TAX1BP1^+^ structures were formed by an accumulation of multiple autophagosomes (**Figure 6B, Fig. S6 and Supplementary videos1-4**). On the other hand, all small TAX1BP1^+^ HSP60^+^ structures were engaged in single cup-shaped LC3B^+^ rings, suggesting they were indeed individual autophagosomes (**Figure 7A**).

**Figure 6:**
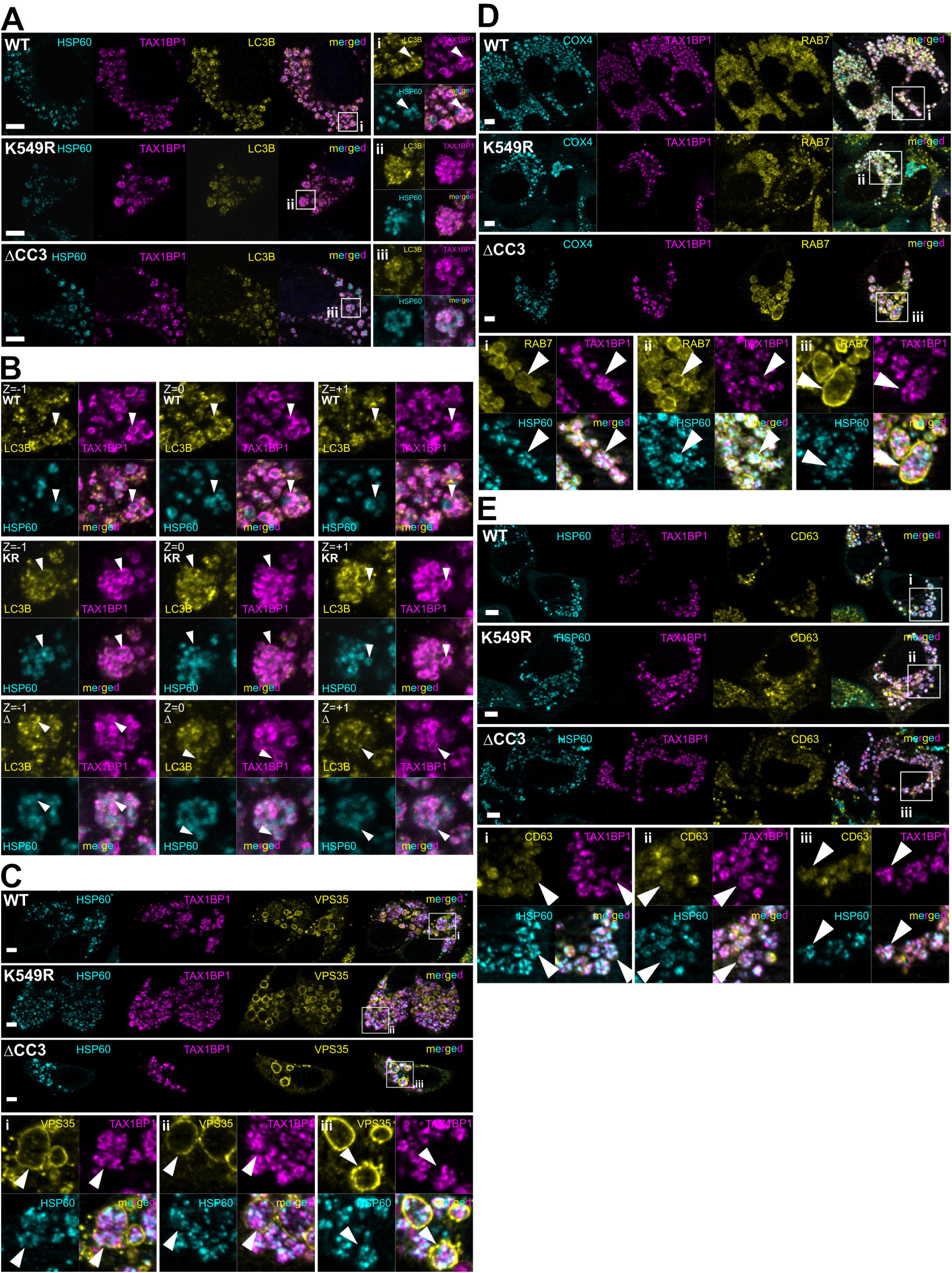
Characterization of enlarged TAX1BP1^+^ vesicles carrying mitochondrial material upon autophagy flux inhibition in 5KO-PARK. (**A**) Representative images of 5KO-PRKN HeLa cells showing enlarged TAX1BP1^+^ HSP60^+^ structures (magenta and cyan, respectively) positive for the autophagosome marker LC3B (yellow) after mitochondria depolarization and BafA for 16 h. Scale bars: 5 μm. (**B**) Z-stack projection of enlarged vesicles identified in (A) for WT, TAX1BP1^K549R^ and TAX1BP1ΔCC3. Thick arrowheads indicate LC3B^+^ TAX1BP1^+^ cup-shaped structures in each Z-plane. Thickness slice: 0.6 μm. (**C-E**) Representative images showing enlarged TAX1BP1^+^ HSP60^+^ structures positive for RAB7 (C), VPS35 (D) and CD63 (E). Zoom insets are shown for WT (i), TAX1BP1^K549R^ (ii) and TAX1BPΔCC3 (iii) for all markers. Scale bars: 5 μm.

**Figure 7:**
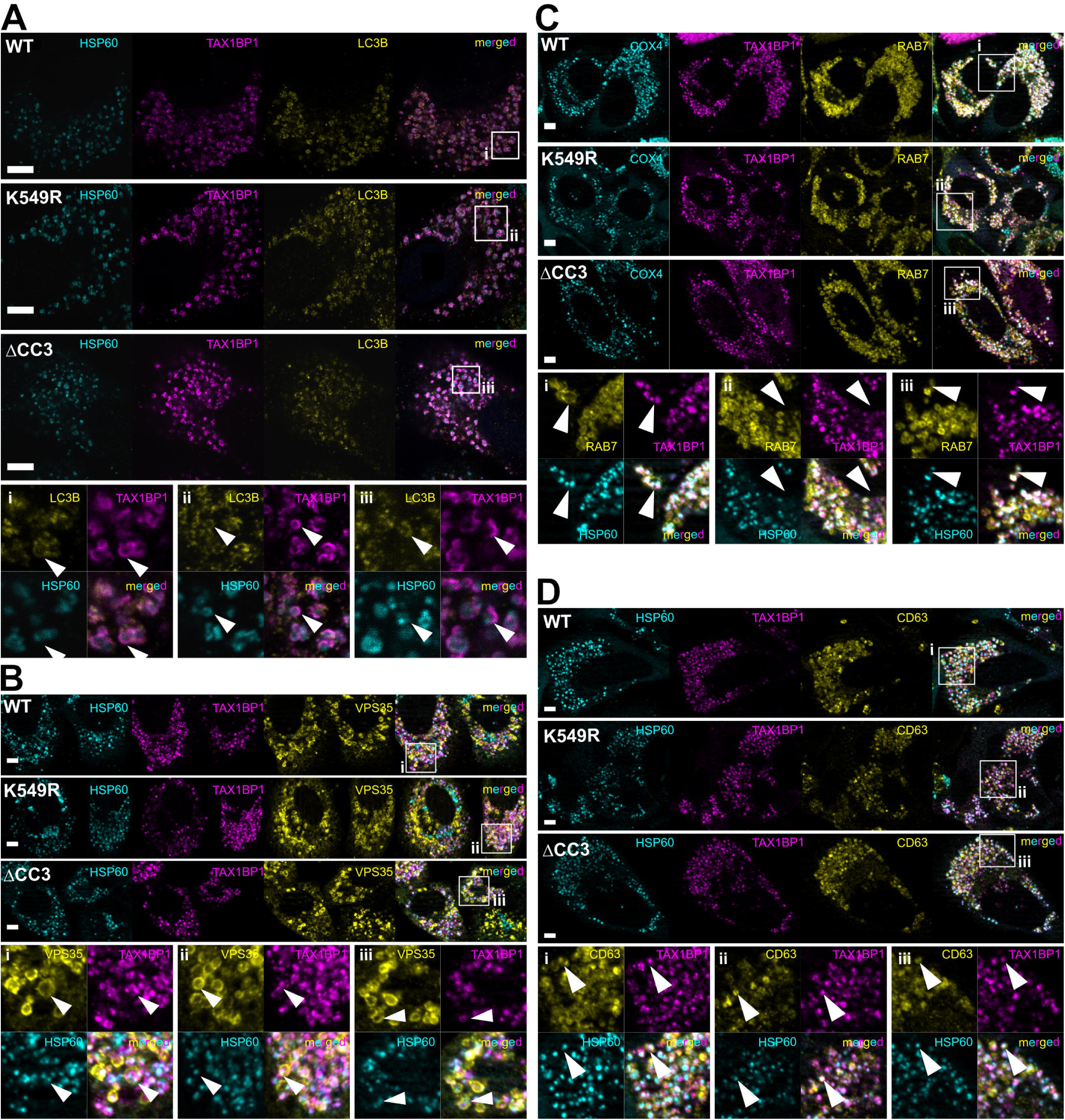
Characterization of small TAX1BP1^+^ vesicles carrying matrix mitochondrial material upon autophagy flux inhibition in 5KO-PARK. (**A-D**) Representative images of 5KO-PRKN HeLa cells showing small TAX1BP1/HSP60-positive structures (magenta, cyan) positive for the indicated markers (yellow): LC3B (A), RAB7 (B), VPS35 (C) and CD63 (D). Zoom insets are shown for WT (i), TAX1BP1^K549R^ (ii) and TAX1BPΔCC3 (iii) for all markers. Scale bars: 5 μm.

Later steps of autophagy-mediated degradation rely on the fusion of sealed autophagosomes with lysosomes [19]. However, prior to autophagosome-lysosome fusion, autophagosome maturation takes place and sealed autophagosomes may additionally fuse with endosomes, generating a new degradative vesicles termed amphisome [22]. Thus, we decided to prove for the late endosomal marker RAB7 (RAB7A, member RAS oncogene family), the member of the retromer complex VPS35 (which is mutated in rare forms of PD) and the late endosomal and marker CD63/LAMP-3 (CD63 antigen or Lysosomal-associated membrane protein 3). Surprisingly, all enlarged TAX1BP1^+^ structures carrying mitochondrial material were surrounded by RAB7 and VPS35 signals, suggesting additional vesicles were engulfing the identified accumulated LC3B^+^ TAX1BP1^+^ HSP60^+^ structures (**Figure 6C and D**). These patterns contrasted with the ones observed for CD63, which rather seemed to be part of the cargo (**Figure 6E**). On the other hand, all regular sized TAX1BP1^+^ structures were positive for all tested endolysosomal markers (**Figure 7B-D),** with the only particularity that the VPS35 and RAB7 rings engulfed single HSP60^+^ and TAX1BP1^+^ structures. These data indicate that the enlarged TAX1BP1^+^ structures are vesicles of endolysosomal nature filled with multiple autophagosomes carrying mitochondrial cargo (amphisomes), while the regular-sized TAX1BP1^+^ structures correspond to individual vesicles or mature autophagosomes.

### Enlarged TAX1BP1ΔCC3^+^ vesicles were also observed in parental HeLa cells with all other complementary autophagy receptors

To confirm the relevance of the previously described phenotype in 5KO-PRKN HeLa cells, similar experiments were performed in WT-Parkin HeLa cells endogenously expressing all other complementary autophagy receptors (NDP52, OPTN, NBR1, SQSTM1/p62, TAX1BP1) (**Fig. S7A**) [11]. Here, siRNA-based transient knockdown experiments were followed by re-transfection of the desired TAX1BP1 constructs and subsequent CCCP-mediated depolarization combined with BafA (**Figure 8A**). Confirmation of efficient silencing was additionally performed via protein expression of total TAX1BP1 (**Figure 8A, lower panel**). Overall, TAX1BP1 puncta in all conditions co-localized with the mitochondrial marker HSP60 (**Figure 8A**). Surprisingly, the mean size of all observed TAX1BP1^+^ puncta per cell significantly increased upon siRNA knockdown and further TAX1BP1 re-transfection in comparison to controls (**Figure 8B**). Indeed, when classifying TAX1BP1 puncta size between small (<50 px^2^), intermediate (≤150 px^2^) or enlarged (> 150 px^2^) (**Fig. S7B and C**), the majority of TAX1BP1^+^ puncta were of small or intermediate sizes, while larger puncta represented a much reduced percentage of the total TAX1BP1^+^ puncta in all conditions (**Fig. S7C**). Of note, there was a 2-fold significant increase of the intracellular percentage of larger TAX1BP1ΔCC3^+^ puncta in comparison to WT TAX1BP1 conditions, reaching 5% of total TAX1BP1ΔCC3^+^ puncta identified (**Figure 8C**). An additional increase of the percentage of larger puncta profiles was observed upon WT TAX1BP1 transfection when compared to transfection and silencing controls, suggesting the overexpression of TAX1BP1 could increase its puncta size (**Figure 8C**). Furthermore, while the number of cells with large TAX1BP1^+^ puncta was slightly higher in TAX1BP1ΔCC3 in comparison to re-transfected WT or TAX1BP1^K549R^, the differences were only significant when compared to siRNA controls (**Figure 8D**). Consistent with previous experiments in 5KO-PRKN Hela, TAX1BP1^+^ enlarged puncta also co-localized with the mitochondrial proteins HSP60 in WT-Parkin HeLa (**Figure 8A**). Next, to relate TAX1BP1 mitochondrial removal efficiency upon expression of the different TAX1BP1 constructs in WT-Parkin HeLa, co-localization analysis of HSP60 and TAX1BP1 was performed. First, the percentage of HSP60 foci co-localizing with TAX1BP1^+^ puncta of all sizes was analyzed (**Figure 8E**). An increased percentage of HSP60 co-localizing with TAX1BP1 was observed upon silencing and TAX1BP1 re-transfection in comparison to controls, with no significant differences among all three TAX1BP1 conditions (**Figure 8E**). These data indicated that all TAX1BP1 constructs enhanced the recognition of HSP60 material, in agreement with previous experiments where overexpression of TAX1BP1 had a positive effect on mitochondrial degradation (**Fig. S2B and C**). Next, HSP60 foci co-localizing with enlarged TAX1BP1^+^ puncta were analyzed (**Figure 8F**). Here, while there was a slight increase of the percentage of HSP60 foci recognized by enlarged TAX1BP1ΔCC3^+^ puncta, no significant differences were found among WT and mutants under silencing conditions (**Figure 8F**). However, it is worth noting that the mean area of the identified HSP60 foci co-localizing with TAX1BP1ΔCC3 and TAX1BP1^K549R^ were larger than the ones observed in WT or control conditions, while no differences were observed for WT compared to controls (**Figure 8G**).

**Figure 8:**
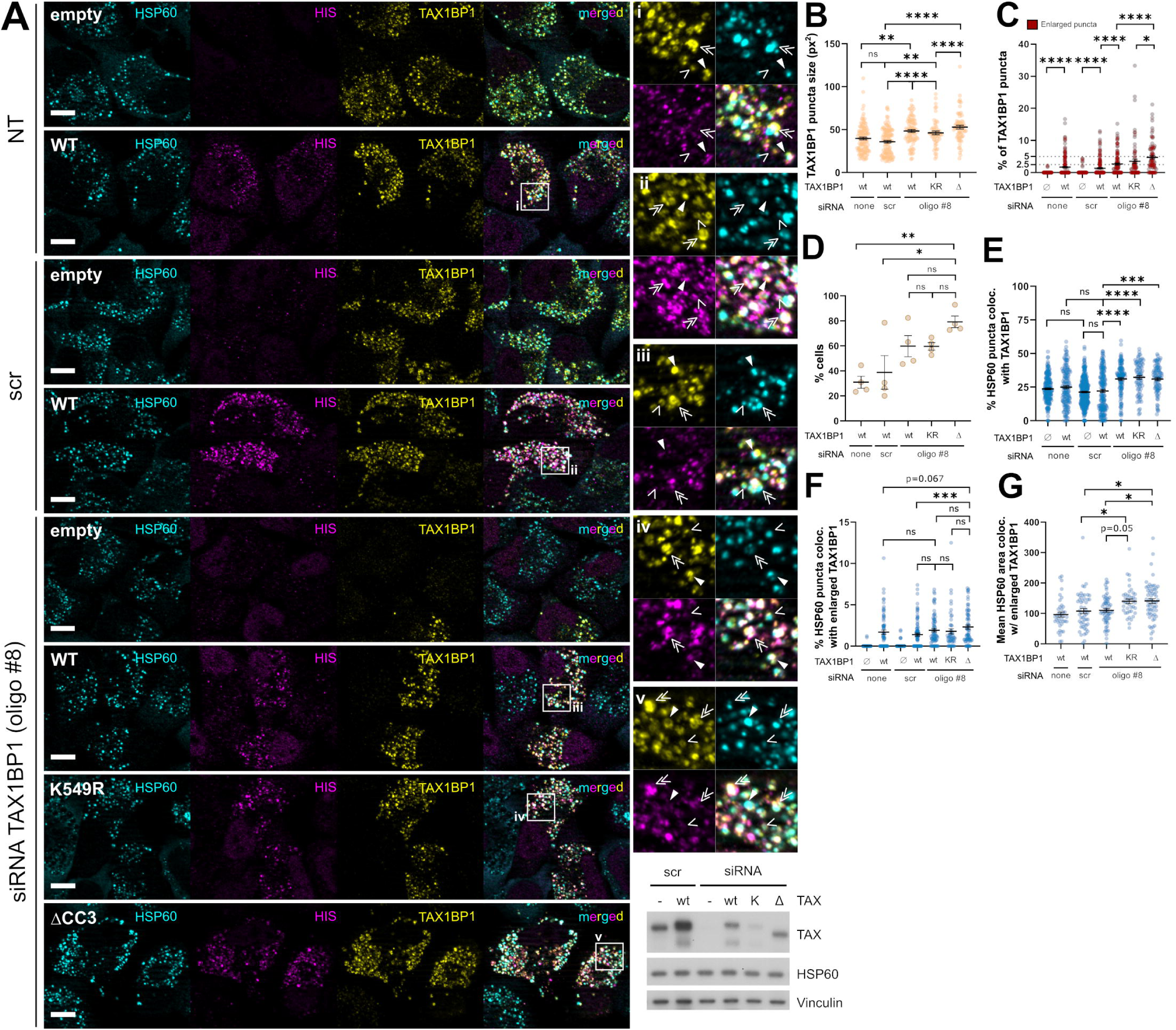
Characterization of TAX1BP1^+^ structures identified in WT-Parkin Hela cells upon autophagy flux inhibition and extended depolarization. (**A**) Representative images upon siRNA mediated knock-down of TAX1BP1 in WT-Parkin HeLa cells showing TAX1BP1^+^ structures (magenta) carrying mitochondrial material (HSP60, cyan). Scale bars: 10 μm. Zoom insets are shown for the indicated conditions (right side, i-v). Lower panel of western blot insets illustrates silencing and re-transfection efficiency (n=2). (**B-G**) CellProfiler based analysis of TAX1BP1 puncta size or HSP60 material colocalizing with TAX1BP1 puncta in WT-Parkin HeLa cells after siRNA mediated TAX1BP1 knockdown and TAX1BP1 re-transfection upon CCCP and BafA treatment. (B) Mean TAX1BP1^+^ puncta size. (C) Percentage of intracellular enlarged TAX1BP1^+^ puncta per cell. (D) Percentage of total cells harboring enlarged TAX1BP1^+^ puncta. (E) Percentage of total HSP60 foci colocalizing with TAX1BP1. (F) Percentage of total HSP60 foci colocalizing with enlarged TAX1BP1^+^ puncta. (G) Mean HSP60 foci area colocalizing with enlarged TAX1BP1^+^ structures.

In short, we observed an increased prevalence of enlarged TAX1BP1ΔCC3 structures capable of recognizing increased amounts of HSP60 in comparison to WT. Similarly, TAX1BP1^K549R^ enlarged structures are able to engage with increased amounts of HSP60, however the prevalence of TAX1BP1^K549R^ enlarged structures is less abundant than TAX1BP1ΔCC3, suggesting a milder phenotype for the site-substitution mutant. Thus, these data indicated that the enlarged TAX1BP1ΔCC3^+^ structures observed in 5KO-PRKN HeLa cells were of dominant nature, despite the partial rescue observed in WT-parkin HeLa, where all complementary autophagy receptors are present.

### Reduced autophagosome formation in TAX1BP1ΔCC3

Next, we hypothesized that the observed accumulation of enlarged amphisomes in TAX1BP1ΔCC3 upon autophagy blockage could be due to defects in autophagy initiation flux at earlier stages. In order to check this, short mitochondria depolarization was induced by CCCP or OA treatment and co-localization of TAX1BP1 and LC3B on mitochondria was further analyzed.

As expected, TAX1BP1-HSP60 co-localization upon CCCP or OA depolarization was significantly increased when compared to control conditions (**Figure 9A and Fig. S8A-B**). However, while all TAX1BP1 mutants were able to translocate to mitochondria upon depolarization, a significant reduction on TAX1BP1-HSP60 overlap was observed for TAX1BP1ΔCC3 while no changes in intensity correlation were detected (**Figure 9B-C**). These data are in agreement with previous experiments where we observed a decrease in TAX1BP1ΔCC3 overlap on COX4^+^ or SSBP1^+^ mitochondrial material (**Figure 4E-F**). Along those lines, mitochondrial area was found increased in TAX1BP1ΔCC3 upon CCCP treatment only (**Fig. S8C-D**), in agreement with previous experiments (**Figure 4B-D**).

**Figure 9:**
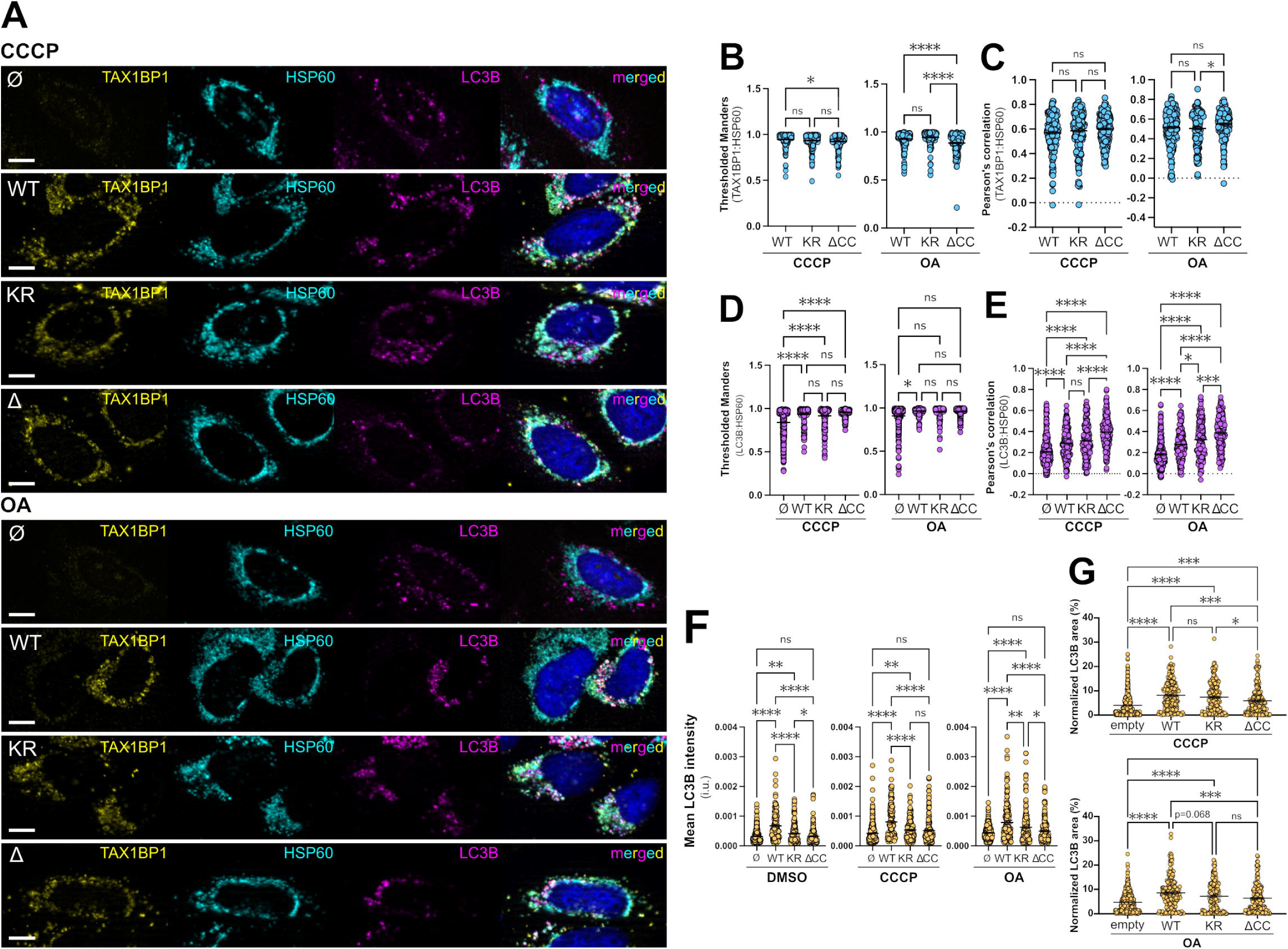
Co-localization analysis of TAX1BP1 and LC3B on depolarized mitochondria upon overexpression of WT, TAX1BP1^K549R^ or TAX1BP1ΔCC3. Depolarization was performed for 4 hours with CCCP or OA, as indicated. (**A)** Representative images showing TAX1BP1 (yellow), LC3B (magenta) and HSP60 (cyan) after 4 h of depolarization. Scale bars: 8 µm. (**B-E**) Colocalization analysis of TAX1BP1-HSP60 (B-C) and LC3B-HSP60 (D-E), represented with thresholded Manders coefficient or Pearson’s correlation coefficient for the respective treatments and TAX1BP1 transfection, as indicated. (**F**) Mean LC3B intensity observed upon the indicated treatments and TAX1BP1 transfection. i.u: intensity units. (**G**) LC3B area relative to total cell area. In all panels, data of individual cells is represented by single dots. All data come from at least three independent experiments. KR: K549R, ΔCC: ΔCC3.

Next, to further understand the potential defects in TAX1BP1ΔCC3-dependent autophagy initiation, LC3B colocalization with HSP60 was analyzed (**Figure 9D-E** and **Fig. S8E**). As expected, LC3B-HSP60 overlap upon CCCP or OA depolarization was significantly increased when compared to control conditions (**Figure S8E**). On the other hand, no changes in LC3B overlap with mitochondria were observed for any of the mutants in comparison to WT TAX1BP1 (**Figure 9D**). However, there were significant changes in LC3B-HSP60 signal correlation for both TAX1BP1 mutants, as a significant increase in LC3B-HSP60 intensity correlation was observed for TAX1BP1ΔCC3 under both CCCP and OA depolarization (**Figure 9E**). Changes in TAX1BP1^K549R^ were only significant under OA treatment conditions (**Figure 9E**). Importantly, this was not a result of increased LC3B intensity in TAX1BP1ΔCC3, as we observed a significant decrease in LC3B intensity for both TAX1BP1 mutants in all conditions (**Figure 9F** and **Fig. S8F**). Similarly, there were also important changes on the LC3B signal area (**Figure 9G**). As expected, TAX1BP1 expression alone induced a significant increase in LC3B area and this was accentuated upon mitochondria depolarization (**Fig. S8G**). However, such increase was not as pronounced in TAX1BP1ΔCC3 after depolarization, as we observed a significant reduction of the total LC3B area in TAX1BP1ΔCC3 when compared to WT TAX1BP1 under both CCCP and OA treatment conditions (**Figure 9G**).

These data strongly suggest that autophagosome formation induced by TAX1BP1ΔCC3 may be compromised due to a reduced recognition of depolarized mitochondria under these conditions, which results in a generally reduced autophagosome production.

### Mass-spectrometry analysis of TAX1BP1ΔCC3 pulldown reveals defects in intracellular trafficking processes

In order to get an idea of the global processes occurring upon TAX1BP1ΔCC3 expression and try to pinpoint specific TAX1BP1 interactors that could be indicative of distinct non-classical autophagic processes, mass-spectrometry analysis of TAX1BP1 precipitates was performed. Here, HIS-tagged TAX1BP1 was co-precipitated under native conditions by means of Ni-NTA pulldown and the pulled down fractions were further analyzed by mass-spectrometry. Both conditions of depolarization only (16 h CCCP) or depolarization combined with autophagic flux inhibition (16 h CCCP + BafA) were analyzed.

Identified protein groups and peptides showed a minor increase for cells expressing WT and mutant TAX1BP1 when compared to the empty plasmid (**Fig. S9A-B**). Furthermore, comparisons of the numbers of protein groups connected to mitochondria as well as the autophagy and vesicular transport machineries revealed no sample specific differences (**Fig. S9A**). The overall sample correlations were high and principal component analysis indicated that the main differences between samples stemmed from variations between replicates (**Fig. S9C-D**). As further quality control of the dataset, the number of identified peptides for TAX1BP1 peptides and protein quantity were investigated. As expected, TAX1BP1 was found to be absent in the control samples (**Fig. S9E-F**). Surprisingly, TAX1BP1^K549R^ quantity values were lower in comparison to WT and TAX1BP1ΔCC3 (**Figure S9F**). Statistical differences were analyzed between TAX1BP1ΔCC3 and WT upon CCCP treatment only (**Fig. S10A and Table S2**) or in combination with BafA (**Figure 10A and Table S1**). Here, when comparing TAX1BP1ΔCC3 with WT after extended depolarization only, an increased co-precipitation of inner mitochondrial proteins was identified (**Fig. S10A**). These findings were in line with the identified delayed mitochondrial degradation phenotype of TAX1BP1ΔCC3 during earlier depolarization (**Figure 4B-D**). On the other hand, when comparing TAX1BP1ΔCC3 and WT upon CCCP combined with BafA inhibition, multiple proteins involved in vesicle-mediated transport, mitophagy or mitochondria localized proteins were significantly regulated (**Figure 10A**).

**Figure 10:**
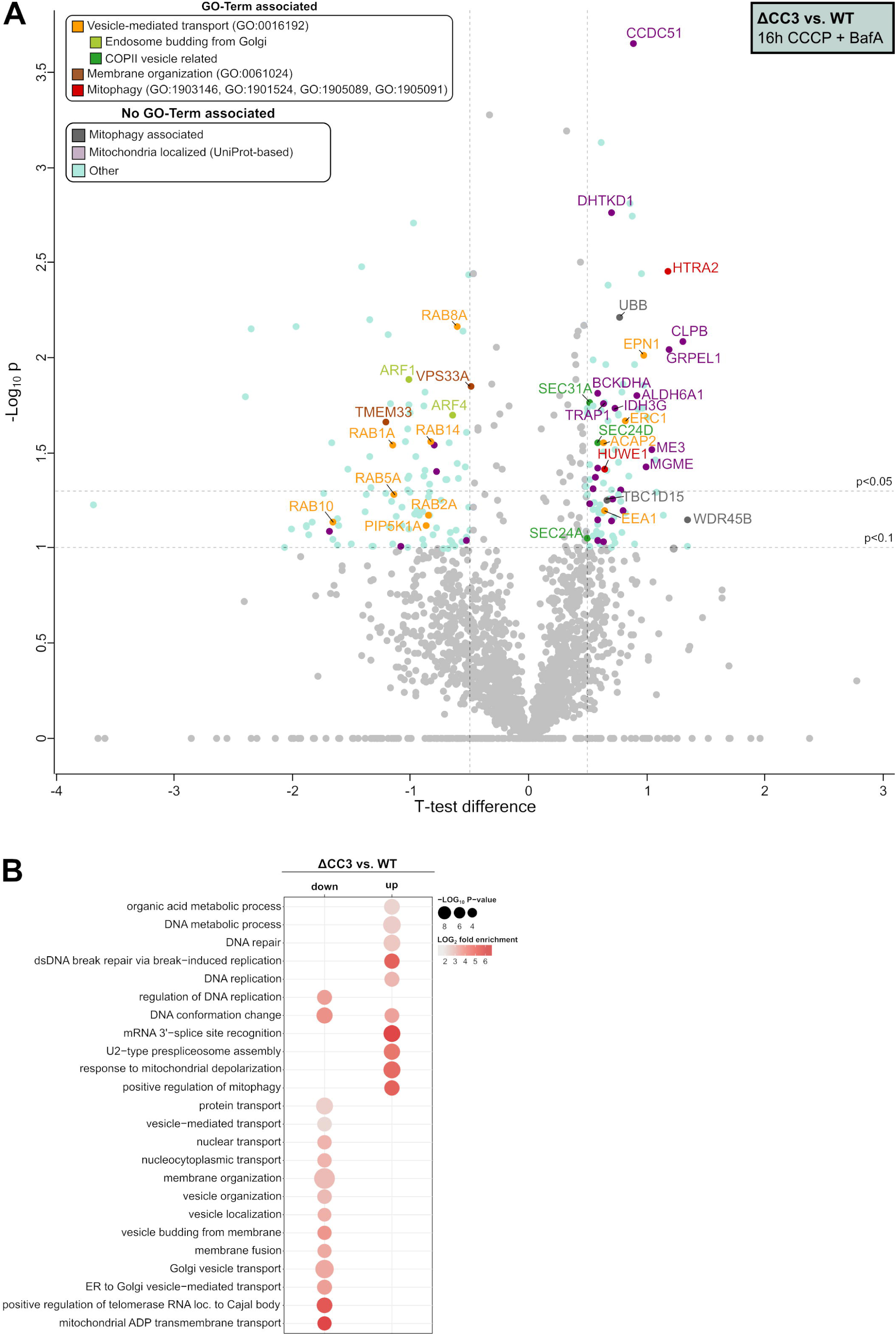
Mass spectrometry analysis of TAX1BP1 native pulldown indicates defects in protein trafficking processes in TAX1BP1ΔCC3. (**A**) Statistical analysis of TAX1BP1ΔCC3 vs. WT TAX1BP1 pulldowns represented by the log_10_ p-value plotted against student’s T-test difference. Significance lines represent p ≤ 0.05 and p ≤ 0.1, as indicated. Proteins are grouped according to the following GO-Terms: vesicle-mediated transport in orange (GO:0016192), membrane organization in brown (GO:0061024) and mitophagy in red (GO:1903146; GO:1901524; GO:1905089; GO:1905091); mitochondria localization in lilac (based on UniProt annotation) or if they are associated with PINK1/parkin mitophagy in grey (based on literature). Proteins related to the GO-Term vesicle-mediated transport (GO:0016192) are classified according to their functionality (based on UniProt and literature); between endosome budding from Golgi (light green) or COPII vesicle related (dark green). (**B**) Overrepresentation analysis of biological processes for significantly regulated proteins by Fisher exact testing.

Significantly up- and downregulated proteins were further analyzed for statistical overrepresentation on GO biological processes for the condition of depolarization combined with autophagy flux blockage only, using the panther database (www.pantherdb.org, version 19.0) [23]. Upon comparing the protein signatures interacting with TAX1BP1ΔCC3 against WT under CCCP and BafA treatment combined, GO-term analysis indicated a downregulation of several vesicle-related biological processes, some of them related to retrograde transport or vesicle budding. Moreover, an upregulation of DNA replication, repair and translation processes and an additional upregulation of mitophagy response processes related to mitochondria depolarization were identified (**Figure 10B**).

When analyzing the protein signatures interacting with TAX1BP1ΔCC3 under these conditions in more detail, a downregulated co-precipitation of TAX1BP1 with several proteins involved in intracellular trafficking was identified (**Figure 10A**). Those included the Rab-GTPases RAB8A, RAB14 and RAB1A, proteins involved in membrane organization (TMEM33 [Transmembrane protein 33], VPS33A [vacuolar protein sorting-associated protein 33A]) and other proteins involved in endosome budding from the Golgi apparatus (ARF1 [ADP-ribosylation factor 1], ARF4 [ADP-ribosylation factor 4]). On the other hand, an upregulated interaction with ubiquitin and inner mitochondrial proteins was found (**Figure 10A**). This later finding correlated with previous data showing an increased amount of enlarged TAX1BP1ΔCC3 positive vesicles carrying increased amounts of mitochondrial material (**Figure 5A and G and Fig. 8G**) as well as with the presence of Ub-TOM20 after short depolarization (**Figure 3B and J**). Additionally, several vesicular-trafficking related proteins were identified to increasingly co-precipitate with TAX1BP1ΔCC3. Those were: three COPII vesicle-related proteins (SEC31A, SEC24D and SEC24A [protein transport protein Sec24A/D/31A]) [24]; the endocytosis-related protein EPN1 (epsin-1) [25]; ERC1 (ELKS/Rab6-intercting/CAST family member 1) known to be involved in exocytosis [26,27]; and the GTPase activating protein ACAP2 (Arf-GAP with coiled-coil, ANK repeat and PH domain-containing protein 2), involved in endosomal trafficking [28] (**Figure 10A**). Moreover, at a lower significance level (p-value<0.1), an upregulation of autophagy-related protein WDR45B/WIPI3 (WD repeat domain phosphoinositide-interacting protein 3) as well as the Rab7 GTPase activator TBC1D15 (TBC1 domain family member 15) were identified (**Figure 10A**), both also implicated in mitophagy or general autophagy [29–32]. Importantly, no significant interaction with ubiquitin or vesicle-related proteins was observed for TAX1BP1^K549R^ while an additional interaction with mitochondrial proteins was identified (**Fig. S10B and Table S1**).

These data indicated that TAX1BP1ΔCC3 under these conditions was capable of dysregulating vesicle-trafficking processes while being able to increasingly interact with ubiquitinated mitochondrial targets. Overall, these findings reinforced the hypothesis that removing the highly ubiquitinatable domain CC3 of TAX1BP1 impaired its mitochondrial targeting efficacy which may additionally have an effect at later stages of autophagy, when autophagosomes are formed and vesicular transport is essential for autophagosome maturation.

## Discussion

Complete mitochondria elimination during PINK1/Parkin mitophagy is subject to autophagy receptor-dependent recognition of ubiquitinated mitochondria. In this context, five autophagy receptors have been linked to PINK1/parkin mitophagy: NBR1, SQSTM1/p62, OPTN, NDP52 and TAX1BP1 [33]. PTMs are known to play a regulatory role on autophagy receptor functionality during selective autophagy [34]. Here we show that, in a context of PINK1/parkin mitophagy, TAX1BP1 ubiquitination at its CC3 domain can regulate the delivery of depolarized mitochondria to either a classical autophagy-dependent degradative pathway or to an alternative pathway that is more dependent on the endolysosomal machinery, without dramatically compromising the elimination of defective mitochondria.

TAX1BP1 ubiquitination at K549 or other lysines within its CC3 domain have been described under similar mitochondria depolarizing conditions as the ones reported in this study (**Figure 1A-B**) [18,35], strongly suggesting a regulatory role for this PTM. Indeed, we could observe that TAX1BP1 ubiquitination did not direct the autophagy receptor for early proteasomal degradation (**Fig. S1A**) but instead, ubiquitinated TAX1BP1 was found enriched in mitochondria fractions upon short depolarization (**Figure 2B**). The limited effects of the K549 point mutation suggests complexity and redundance in the TAX1BP1 ubiquitination pattern. This study relies on the deletion of the whole CC3 domain to understand the role of TAX1BP1 ubiquitination and the data presented here indicate that the CC3 deletion does not impair the overall functionality of the autophagy adaptor. However, we cannot discard that the entire CC3 domain deletion subtly perturbs TAX1BP1 structure, accounting for potential effects beyond ubiquitination. Nevertheless, the CC3 domain is richer in lysins in comparison to the other coiled-coiled regions of TAX1BP1, and the lysines within the CC3 domain are highly conserved across mammalian species (**Fig. S3**). Indeed, when deleting the CC3 domain of TAX1BP1, a clear decrease in total TAX1BP1 ubiquitination was observed, both after short (4 h CCCP) and extended depolarization combined with autophagy flux inhibition (16 h CCCP + BafA) (**Figure 3B-E**, **Figure 5E and Fig. S2C**). Interestingly, when checking TAX1BP1 predicted structure in AlphaFold [36], the structure of the regions connecting the different CC domains and the ubiquitin binding domain (UBD) with the CC3 are of lowest confidence (**Figure 11A**). This could be indicative of a highly dynamic behavior of those regions, which could in turn influence the orientation of the distinct CC and UBD regions. Thus, while there is no evidence for a potential structural interaction between the CC3 and UBD regions based on the AlphaFold model, it is tempting to speculate that the ubiquitination of multiple lysines at the CC3 could influence the orientation or availability of the UBD region (**Figure 11B**).

**Figure 11:**
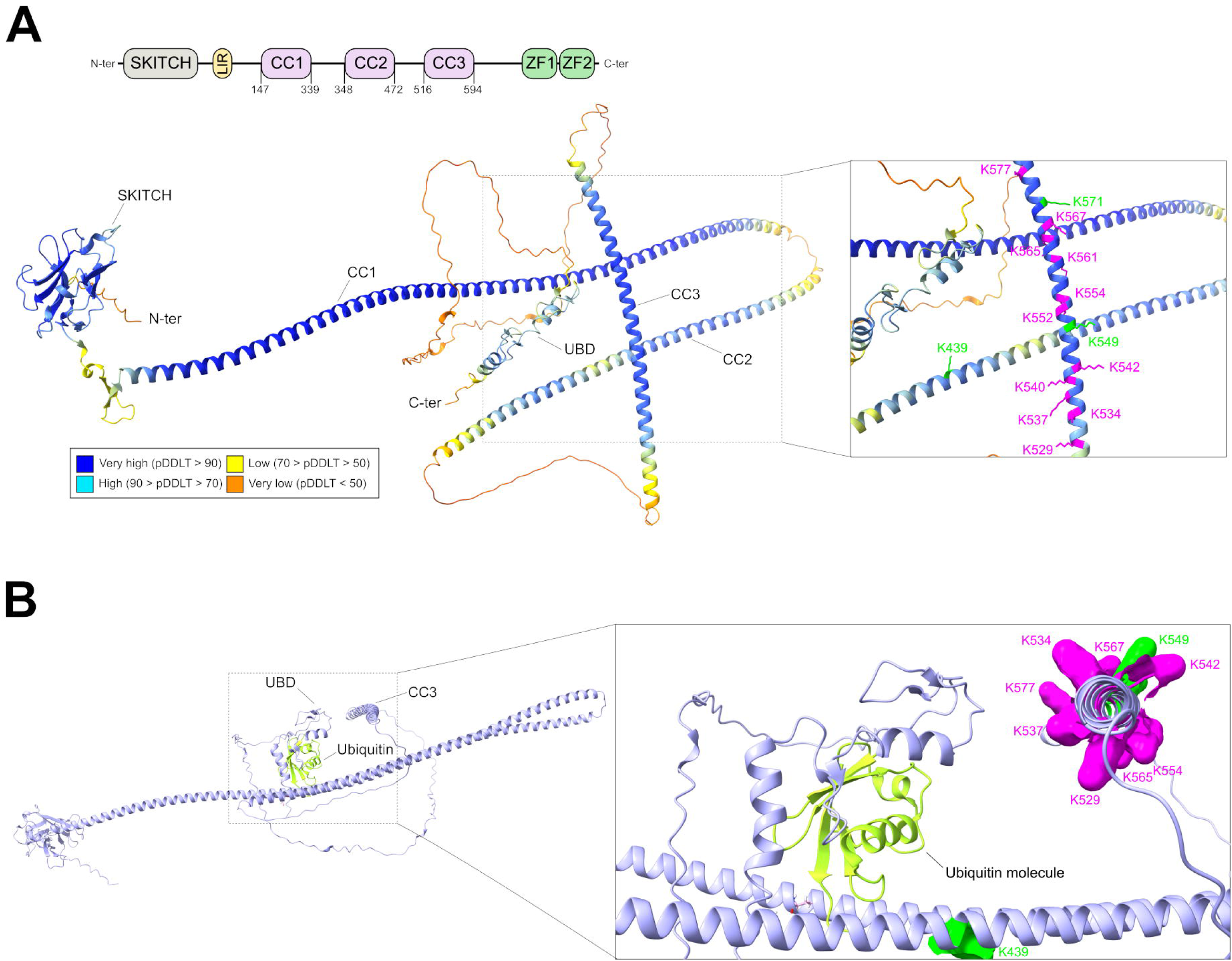
Structural visualization of TAX1BP1 using the predicted Alphafold structure. (**A**) Domain organization of the coiled-coil domains of TAX1BP1 according to Alphafold (top panel). Predicted Alphafold structure of TAX1BP1 with the indicated per-residue model confidence scores (pLDDT). In the zoom panel, identified lysines in the mass-spectrometry screen are highlighted in green. Highlighted in magenta are the remaining lysines of the CC3, which were not identified via MS. (**B**) TAX1BP1, structure in combination with a ubiquitin molecule at its ubiquitin-binding domain (UBD). Zoom panel shows lysines of CC3 (residue surface) in close proximity to the UBD. Highlighted lysines of CC3 are shown in green, if they were identified in the mass-spectrometry screen. The rest of lysines from CC3 are shown in magenta. Images created with ChimeraX [63].

Abolishing TAX1BP1 ubiquitination at CC3 domain by deleting its domain did not impair the degradation of OMM proteins but had a negative impact in the targeting of Ub-TOM20 (**Figure 3B, F-J**) and promoted a slower clustering of mitochondria after short depolarization (**Figure 4A-D**). Moreover, while CC3 ubiquitination was proven not essential for TAX1BP1 translocation to mitochondria (**Figure 4A**), a lack of it did significantly reduce the distribution of the autophagy receptor on CCCP- and OA-depolarized mitochondria (**Figure 4E-F and Figure 9A-B**). Thus, suggesting that the delayed mitochondrial degradation and clustering of TAX1BP1ΔCC3 was possibly due to an impairment on depolarized mitochondrial recognition. One of the first reports describing TAX1BP1 domain structure and its first links to immunity-related functions, described three different coiled-coil domains localized between the now known SKITCH and UBA domains [37]. Interestingly, Ling and Goeddel identified the CC1 and CC2 domains to be essential for TAX1BP1 homodimerization under basal conditions, while the region flanking the CC3 domain seemed to play a minor role in its assembly [37]. Similarly to TAX1BP1, the autophagy receptor NDP52 also possesses a central CC region recently identified to be crucial for NDP52 homodimerization in the context of mitophagy [38,39]. Based on this, it is tempting to speculate that the differences observed in TAX1BP1ΔCC3 distribution on mitochondrial material might be due to a reduced homodimerization capacities. However, given that the recognition of depolarized mitochondria was significantly reduced in TAX1BP1ΔCC3 (**Figure 4E-F and 9A-B**) and this phenomenon was accompanied by a reduced formation of LC3B^+^ vesicles, as indicated by a reduced LC3B intensity and total LC3B area in TAX1BP1ΔCC3 (**Figure 9F-G**), it seems possible that the delivery of defective mitochondria in TAX1BP1ΔCC3 could be compromised or delayed from this step on. Along these lines, our study also reveals that, even though the lack of ubiquitination at CC3 does not impair complete mitochondrial degradation (**Figure 5C**), the degradative pathways governing this process may differ between WT and TAX1BP1ΔCC3, as an increased presence of enlarged degradation vesicles was observed in TAX1BP1ΔCC3 only (**Figure 5 and Fig. S5**), even under conditions where all other autophagy receptors were present (**Figure 8 and Fig. S7C**). It is noteworthy that the here identified enlarged TAX1BP1^+^ vesicles were prominently surrounded by the PD gene product VPS35. These findings may suggest a functional link between the mitophagy pathway regulated by the most common recessive gene products PINK1 and parkin together with the rare VPS35 gene product, suggested to be involved in vesicle trafficking effects of the common dominant PD gene product LRRK2 (Leucine-rich repeat serine/threonine-protein kinase 2). While this latter connection in our work may be quite superficial, recent work by the group of Vandenberghe W. and colleagues has made evident a remarkable functional link between VPS35-LRKK2 in the context of PINK1/parkin mitophagy [40]. The authors identify the novel VPS35 mutation p.D620N to not compromise the initiation steps of PINK1/parkin mitophagy but significantly impair the recruitment of the autophagy receptor OPTN on depolarized mitochondria [40]. While at this point it still remains yet to be elucidated if TAX1BP1 ubiquitination may have any impact on this molecular axis, our mass-spectrometry data seems to confirm defects in vesicular trafficking, as many targets related to vesicle-mediated transport were downregulated in TAX1BP1ΔCC3 (**Figure 10A**). Nevertheless, we could not identify a unique endolysosomal marker specific for the enlarged degradative vesicles only under these conditions, as all enlarged and small vesicles were positive for the same markers (LC3B, VPS35, RAB7 and CD63) (**Figure 6 and Figure 7**). However, it is important to emphasize that strong structural differences were clearly observed between small and enlarged TAX1BP1^+^ degradative vesicles, being the enlarged ones more prevalent in TAX1BP1ΔCC3 than in WT. More precisely, the enlarged TAX1BP1ΔCC3^+^ structures contained numerous TAX1BP1^+^ LC3B^+^ vesicles carrying mitochondrial material (**Figure 6A-B, Fig. S6 and Supplementary videos 1-4**), where globally surrounded by RAB7^+^ and VPS35^+^ vesicles (**Figure 6C-D**) and were additionally positive for CD63 (**Figure 6E**). On the other hand, small vesicles found in all TAX1BP1 constructs consisted of single TAX1BP1^+^ LC3B^+^ positive vesicles also carrying mitochondrial material and were equally positive for all the tested endolysosomal markers (RAB7, CD63 and VPS35) (**Figure 7**). These data seem to indicate that the degradation pathways involved in WT and ubiquitination deprived TAX1BP1 are possibly of the same nature and the identified enlarged and small vesicles are the result of autophagosomes fused with endosomes (amphisomes) or lysosomes (autolysosomes) [22], potentially indicating that the complex enlarged TAX1BP1 structures might be the result of defects in autophagosome maturation. Autophagosome fusion with endosomes is known to be necessary for autophagosome maturation as it increases the pH of the degradative vesicle and is considered a previous step for lysosome fusion [22]. On the other hand, autophagy flux inhibition via BafA does not specifically hinder the fusion of autophagosomes with lysosomes [41], thus we cannot discard that the observed enlarged TAX1BP1 positive vesicles simply correspond to autolysosomes with more complex LC3B-TAX1BP1-HSP60 cargo. It is important to note that TAX1BP1 has been reported to interact with the cytoskeletal motor MYO6 (unconventional myosin-VI), known to be involved in endocytic transport and to facilitate autophagosome maturation by interacting with the UBD of TAX1BP1 [42]. Interestingly, compromising autophagosome maturation via transient knock-down of MYO6, induced the formation of immature enlarged autophagosomes positive for SQSTM1/p62 [42]. Another recent example of the relationship between degradative vesicle size and endosome/autopagosome maturation was achieved by transient knockdown of individual HOPS complex subunits, known to play an essential role in endosome and autophagosome fusion with lysosomes [43]. Based on this, it seems reasonable to think that the enlarged vesicles observed in TAX1BP1ΔCC3 are the result of impaired autophagosome maturation driven by TAX1BP1ΔCC3 alone and further accentuated by BafA treatment.

Additionally, it is important to emphasize that the formation of enlarged degradative vesicles was not only observed in 5KO HeLa (**Figure 5B-D, F-H and Figure 6**) but also in parental WT-Parkin HeLa with all complementary autophagy receptors (**Figure 8**). In this case, the observed TAX1BP1ΔCC3 positive structures were generally smaller as the ones observed in 5KO HeLa. However, amongst the regularly sized TAX1BP1 positive structures, an increased proportion of enlarged TAX1BP1ΔCC3 positive structures was observed (**Figure 8C-D**). These data indicate that while the presence of complementary autophagy receptors can partially compensate for the observed phenotype, the TAX1BP1ΔCC3 dependent formation of enlarged degradative vesicles can be considered of a dominant nature. Moreover, the mitochondrial material recognized by the TAX1BP1^+^ enlarged vesicles was significantly increased in TAX1BP1ΔCC3 and TAX1BP1^K549R^ in comparison to WT (**Figure 8G**), strongly suggesting a decreased efficiency of ΔCC3, also under conditions of complementary autophagy receptor expression. Thus, as larger TAX1BP1 vesicles were observed in 5KO HeLa when compared to the ones seen in WT-Parkin HeLa, it seems reasonable to think that the expression of complementary autophagy receptors can partially compensate for the TAX1BP1-dependent formation of enlarged degradative vesicles. This is in agreement with previous reports indicating TAX1BP1 as the least efficient autophagy receptor to promote PINK1/parkin mitophagy in 5KO HeLa [11]. In addition, recent reports using proximity-labelling approaches have indicated that TAX1BP1 can interact with other autophagy receptors during mitophagy [44,45] suggesting the cooperation with more efficient autophagy receptors could partially compensate for the observed phenotype in 5KO HeLa.

Upon general GO-Term analysis of TAX1BP1ΔCC3 native pulldown (**Figure 10B**), a significant decrease in interactions related to general vesicle trafficking was identified and is in agreement with previous results showing an increased presence of TAX1BP1ΔCC3 enlarged vesicles under these conditions (**Figure 5B-D, F-H, Figure 6 and Fig. S5**). An upregulated interaction with inner mitochondrial proteins and ubiquitin confirmed a decreased efficiency of ΔCC3 in mitochondria degradation (**Figure 3C**, **Figure 4B-D and Figure 10A**). However, expected TAX1BP1 interactors like RB1CC1/FIP200 (RB1-inducible coiled-coil protein 1) or MYO6 could not be detected for neither of the mutants or WT TAX1BP1. The increased interaction of TAX1BP1ΔCC3 with ubiquitin and the lack of MYO6 co-precipitation may be indicative of delayed ubiquitinated mitochondria recognition, as MYO6 binding is known to compete with ubiquitin at the UBD of TAX1BP1 [42]. On the other hand, an upregulated interaction of TAX1BP1ΔCC3 with the components of COPII vesicle coat complex SEC24A, SEC24D, SEC31A and the receptor-mediated endocytosis EPN1 were identified [46–48] (**Figure 10A**). Similarly, albeit at a lower-significance level, an increased interaction of two autophagy-related proteins involved in autophagosome biogenesis (WDR45B and TBC1D15) was observed (**Figure 10A**). Importantly, COPII vesicles are known to be important source of membrane lipids for the formation of autophagosomes and indeed, recent reports have demonstrated that both SEC24 and SEC31 can interact with ATG9 vesicles, directly influencing autophagosome biogenesis [49–51]. Similarly, the Rab GTPase activating protein TBC1D15 is closely implicated with mitophagy progression via its capacity to modulate active RAB7 levels at the OMM in favor of autophagosome biogenesis [29,30]. On the other hand, WDR45B/WIPI3 is known to localize at nascent autophagosomes as well as in late endosomes and thus seems to also play a role in autophagosomes biogenesis and maturation [31,32]. Thus, given that TAX1BP1ΔCC3 induced (i) a delayed delivery of Ub-TOM20 (**Figure 3J**), (ii) a decrease in mitochondrial clustering as well as a reduced recognition of defective mitochondria (**Figure 4B-F and Figure 9B**) and (iii) promoted a reduced sequestration of LC3B^+^ vesicles upon short depolarization (**Figure 9G**), it seems reasonable to think that the interaction of TAX1BP1ΔCC3 with the autophagosome biogenesis-related proteins at such later stage of the pathway is probably due to a slower capability of the autophagy receptor to recruit autophagy-related proteins in charge of autophagosome biogenesis. Indeed, decreased interactions of TAX1BP1ΔCC3 were observed for the Rab GTPases RAB14, RAB1A and RAB8A, the ER-transmembrane protein TMEM33 and the HOPS complex member VPS33 (**Figure 10A**). Rab GTPases are generally involved in endosomal vesicle trafficking but have also been described to play a role in modulating autophagy: RAB1A has been described to be essential for autophagosome formation [52,53] while RAB14 is implicated in autophagosome maturation [53]. Similarly, VPS33A acts as a tether between autophagosomes and lysosomes to facilitate their fusion [22,54]. Potential defects in autophagosome maturation would be in agreement with the identified sustained ubiquitin interaction, considering that MYO6 interaction competes with ubiquitin at the UBD of TAX1BP1 [42]. Overall, our data suggest that the vesicle-trafficking defects identified in TAX1BP1ΔCC3 may be due to impaired autophagosome biogenesis or maturation and likely derive from a reduced recognition of depolarized mitochondria and a reduced sequestration of LC3B^+^ vesicles at earlier stages of mitophagy (**Figure 12**).

**Figure 12:**
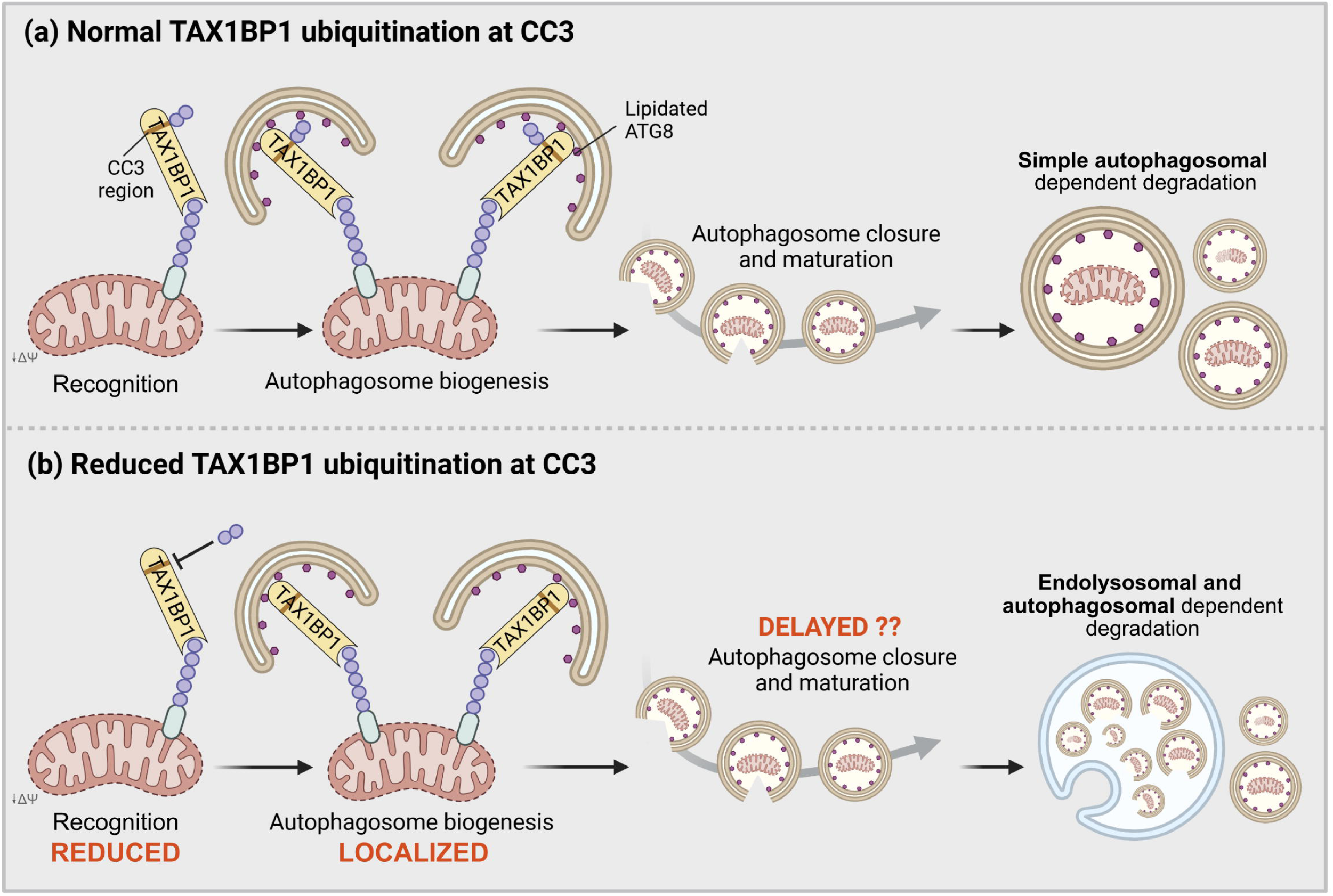
Graphical summary highlighting how reduced ubiquitination on TAX1BP1 can influence the degradative pathway responsible for eliminating defective mitochondria in a PINK1/parkin context. (**A**) Upon PINK1/parkin activation, TAX1BP1 is ubiquitinated at the CC3 domain and translocates to mitochondria. Recognition of ubiquitinated mitochondria via TAX1BP1 promotes autophagosomal delivery. Simple autophagosomal-dependent degradation occurs after successful engulfment of defective mitochondria, autophagosome closure and autophagosome maturation, fostering the degradation of mitochondria. (**B**) Reduced TAX1BP1 ubiquitination or lack of the ubiquitinatable CC3 domain of TAX1BP1 in a context of PINK1/parkin mitophagy follows a slightly different behavior. Here, while TAX1BP1 is able to recognize ubiquitinated mitochondria, its translocation to mitochondria is hampered when compared to WT/normal conditions. This recognition is able to induce the formation of autophagosomes around defective mitochondria in this case, albeit at lower level. Recruitment of LC3-positive vesicles occurs in a more localized manner, which deviates mitophagy dynamics from this point on. Although this pathway leads to successful degradation of defective mitochondria, in this case the prevailed degradative pathway is more dependent on the endolysosomal network rather than on the classical autophagy machinery.

## Conclusions

In this study, we identified the CC3 of TAX1BP1 as a highly ubiquitinated region during PINK1/parkin mitophagy which is not essential for complete mitochondria degradation. However, while reducing CC3 ubiquitination did not really have an impact on OMM protein degradation, other important defects related to mitophagy progression were identified: (i) reduced clearance of ubiquitinated TOM20, (ii) reduced mitochondrial clustering and (iii) reduced TAX1BP1 recognition of depolarized mitochondria, which translated in a (iv) localized presence of LC3B^+^ vesicles on depolarized mitochondria. Blocking autophagy under these conditions had a significant impact on vesicular trafficking, as evidenced by the complex architecture of enlarged TAX1BP1^+^ degradative vesicles which were highly prevalent in the absence of the ubiquitinatable CC3 domain. Our data strongly suggest that the identified vesicular trafficking defects may be driven by a delayed initiation of autophagosome biogenesis as well as an impairment with autophagosome maturation and are a reflection of the identified functional changes of TAX1BP1ΔCC3, which relies on an alternative degradation pathway that is substantially more dependent on the endolysosomal machinery (**Figure 12**).

## Limitations of the study

This study provides evidence for an influence of TAX1BP1 ubiquitination on PINK1/parkin-mediated mitophagy in an idealized, designed HeLa cell culture model. To expand the relevance of the findings provided in this manuscript, more physiological cell models should be investigated, which would require sophisticated manipulations to study if and how TAX1BP1 ubiquitination influences the clearance pathways of defective mitochondria. Given that all the experiments carried out in this study relied on the overexpression of parkin or TAX1BP1 (WT or mutants), future studies carried out for validation should also pay attention on physiologically relevant expression levels of both the E3 ligase and the autophagy adaptor.

## Materials and methods

### Cell culture, treatments and transfections

All cell lines were grown at 37°C and 5% CO_2_ with high-glucose Dulbecco’s modified eagle medium (DMEM). WT-Parkin and 5KO-PRKN HeLa were supplemented with 0.7 mg/ml geneticin (Gibco, 10131-027) to ensure stable parkin expression. HeLa cells stably expressing WT or inactive mutant (C431A) were previously described [55]. To obtain 5KO HeLa stably expressing WT-Parkin (5KO-PRKN HeLa), cells were transfected with BglII linearized pcDNA3.1-3xFlag-Parkin for 24 hours and were further selected with 1.8 mg/ml of Geneticin for 13 days. Upon selection, clones were screened for homogenous Parkin expression. Mitophagy induction was achieved with 10 µM CCCP (Sigma, 555-60-2) for the indicated time points. Proteasome and autophagy inhibition were induced with 10 µM MG132 (Sigma, C2211) or 400 nM BafA (Chemcruz, sc-201550A), respectively. Inhibition treatments were initiated half an hour before CCCP treatment and were maintained in combination with CCCP for the indicated periods. Long CCCP treatment of 5KO-PRKN HeLa was combined with 60 µM Q-VD-Oph (QVD, Selleckchem, S7311) to prevent cell death. Transient transfection of the indicated constructs was performed with FuGENE6® (Promega, E2692), following manufacturer’s instructions. HeLa cells were transfected with 5 nM of siRNA for three consecutive days using Hiperfect (Qiagen, 301709). The following siRNA were used: negative control (AllStars negative, Qiagen, SI03650318); TAX1BP1_7, 5’-CAGATCAATCAGCTAATAATA-3’ (Qiagen, SI02781268); TAX1BP1_9, 5’-AAGGGTCTTACTGAAGTAACA-3’ (Qiagen, SI03036124); TAX1BP1_8, 5’-CAGTCTTTGGCTTATCAATAA-3’ (Qiagen, SI02781296).

### Expression constructs

The construct pcDNA3.1-3xFlag-Parkin has been described previously [56] and was used to transiently express parkin in 5KO HeLa and to generate the stable 5KO-PRKN as previously explained. Full length WT, TAX1BP1^K549R^ and TAX1BP1ΔCC3 cDNA were amplified from commercial vectors obtained from Origene (WT TAX1BP1, RC208233) or eZyvec on demand (K549R and ΔCC3, Quote: D2022-10) and were sub-cloned into pCMV-6HIS vector to ensure equal expression conditions. Obtained constructs were further sequenced prior to use.

### Western blotting

Upon treatment, cells were washed once with phosphate-buffered saline (PBS) and directly harvested. For whole cell lysate samples, cell pellets were resuspended in urea lysis buffer (10 mM Tris, pH 8.0, 100 mM NaH_2_PO_4_, and 8 M urea) after a short centrifugation step (9727 x g for 1 minute [min] at 4°C). Lysates were passed through a 26-gauge needle to break the DNA for at least 5 times. Whole cell lysate fractions were obtained after pelleting debris at 21885 x g for 15 min at 4°C. Protein concentration was measured using the Bradford Protein Assay Dye Reagent Concentrate (Bio-Rad, 5000006) following manufacturer’s instructions. Proteins were separated using 4-12% NuPage (Invitrogen, NP0329BOX), 3-8% Tris-Acetate pre-casted (Invitrogen, EA0378BOX) or self-casted polyacrylamide gels and were run at a fixed voltage of 80-100 V for at least 1,5 hours at room temperature (RT). Proteins were transferred to polyvinylidene fluoride (PVDF) membranes (Merck, IPVH00010) at constant voltage of 25 V for an overnight run or at 100 V for 2 hours, respectively. Upon transfer, membranes were blocked with 5% non-fat dried milk or 5% bovine serum albumin in Tris-buffered saline containing 0.1% Tween-20 (TBS-T). Membranes were incubated with primary antibodies for 2 h at RT or overnight at 4°C. After three consecutive TBT-T washes of 5 min, membranes were incubated with the corresponding secondary antibodies diluted in 5% of non-fat dried milk for 1 hour at RT. Upon additional three TBT-T washes of 5 min, protein bands were visualized using chemiluminescent-horseradish peroxidase (HRP) substrate (Millipore; WBKLS0500) and Ultracruz autoradiography films (Santa Cruz Biotechnology; sc-201697). The following primary antibodies were used: Actin (1:50000, Sigma Aldrich, A5441), Calnexin (1:10000, Enzo Life Sciences, ADI-SPA-860), CS (1:6000, GeneTex, GTX110624), Flag-tag (1:10000, Sigma-Aldrich, F1804), LAMP1 (1:10000, DSHB Hybridoma bank, Clone H4A3), LC3B (1:10000, GeneTex, GT3612), LC3B (1:10000, Cell signaling, 835065), MFN1 (1:10000, Abnova, H00055669-M04), NBR1 (1:10000, ProteinTech, 16004-1-AP), NDP52 (1:10000, ProteinTech, 12229-1-AP), OPTN (1:10000, Atlas antibodies, HPA003360), SQSTM1/p62 (1:10000, BD Biosciences, 610832), Parkin (1:50000, Cell signaling, 4211S), phospho-Ubiquitin (Ser65) (1:1000, Cell Signaling, 52802), TAX1BP1 (1:1000, Novus Biologicals, NBP1-86662), TOM20 (1:30000, Santa Cruz, sc-17764), TOM20 (1:6000, Proteintech, 11802-1-AP), Total Ubiquitin (1:10000, Millipore, MAB1510), VDAC (1:10000, ProteinTech, 66345-1), VDAC (1:30000, Millipore, AB10527), Vinculin (1:50000, Sigma Aldrich, V9131). The following secondary antibodies: Anti-mouse HRP (1:10000, Amersham Pharmacia, 115-035-003), Anti-rabbit HRP secondray (1:10000, Amersham Pharmacia, 111-035-003). Quantification of relative protein levels was obtained via densitometry using ImageJ software. Briefly, individual protein bands were normalized to Vinculin or Actin protein levels.

### Immunofluorescence

HeLa cells were seeded in 0,1 mg/ml PDL (Sigma-Aldrich, P6407) pre-coated round coverslips (Ø: 12 mm). After treatment, cells were fixed with 4% PFA for 20 min. After fixation and three consecutive washes with PBS, permeabilization was performed with 1% Triton-X for 5 min at room temperature (RT). After three washing steps with PBS, blocking was performed at RT for 1 h with 10% fetal bovine serum or donkey serum (blocking buffer). Afterwards, coverslips were incubated with the corresponding primary antibodies diluted in blocking buffer for 2 h at RT. After three washing steps with PBS, coverslips were incubated with secondary antibodies diluted in blocking buffer for 1 h at RT. Following three additional washing steps with PBS, nuclei were stained with 2 µg/ml of Hoechst 33342 in PBS for 5 min at RT. After incubation, three additional washing steps with PBS were performed. Coverslips were finally mounted with fluorescent mounting media (Agligent Dako, S3023). Imaging was performed using an AxioImager microscope equipped with Apotome imaging system (Carl Zeiss, Germany). Images were processed with the ZEN 2.3 lite software and Image J. For LC3B immunostaining, cells were fixed with ice-cold methanol for 2-3 min. Further immunostaining steps were performed as previously described. Image acquisition of endogenous LC3B was realized with a Nikon Ti2e spinning disk confocal microscope. In this case, images were processed using the NIS-Elements software (Nikon, Germany). The following primary antibodies were used: CD63 (1:250, Antibodies online, ABIN94214), COX4 (1:10000, Cell signaling, 4850), HIS-tag (1:500, Addgene, 184180), HSP60 (1:1000, Novus Biologicals, NBP3-05536), LC3B (1:500, Cell signaling, 835065), Parkin(1:10000, Cell signaling, 4211S), RAB7 (1:200, Cell signaling, 95746), TAX1BP1 (1:500, Novus Biologicals, NBP1-86662), VPS35 (1:250, Novus Biologicals, NB100-1397). The following secondary antibodies were used: Donkey anti-sheep Alexa Fluor 647 (1:1000, Invitrogen, A-21448), Donkey anti-mouse Alexa Fluor 488 (1:1000, Invitrogen, A-21202), Donkey anti-mouse Alexa Fluor 568 (1:1000, Invitrogen, A-10037), Donkey anti-rabbit Alexa Fluor 488 (1:1000, Invitrogen, A-21206), Donkey anti-rabbit Alexa Fluor 568 (1:1000, Invitrogen, A-10042), Donkey anti-chicken Alexa Fluor 647 (1:1000, Invitrogen, A78952), Donkey anti-goat Alexa Fluor 648 (1:1000, Invitrogen, A21447), Donkey anti-goat Alexa Fluor 488 (1:1000, Invitrogen, A11055).

Eye-based quantification of TAX1BP1 colocalization on COX4 was performed by taking into account only TAX1BP1^+^ cells with COX4 clumps after 4 h CCCP treatment, to account for parkin-positive cells. TAX1BP1 behavior was classified between complete or localized apposition of COX4 clump depending on its distribution. In situations where both TAX1BP1 behaviors were observed in the same TAX1BP1^+^ cell, localized apposition of TAX1BP1 with COX4 cluster was considered. Quantification of complete mitochondria elimination upon TAX1BP1 transfection was achieved by only taking into account parkin-positive cells (empty control) or cells positive for both parkin and TAX1BP1 (TAX1BP1 transfection) after CCCP treatment. Cells were considered to be cleared of mitochondria where no SSBP1 puncta were observed. Quantification of small or enlarged TAX1BP1 positive structures after extended depolarization combined with BafA inhibition was performed by taking into account only parkin-positive cells. Cells were classified between containing small or enlarged TAX1BP1 structures according size. In situations where both enlarged and small TAX1BP1 structures were observed in the same TAX1BP1^+^ cell, the cell was quantified as containing enlarged TAX1BP1 structures.

### Mitochondria fractionation

Mitochondria purification was performed as described in [57] with some modifications. Cell pellets were first resuspended in mitochondrial buffer (MB) (210 mM mannitol, 70 mM sucrose, 10 mM HEPES, 1 mM EDTA at pH 7.5) and further cell lysis was performed mechanically using a glass homogenizer. After a first lysing series of 30 mechanical stokes, cell homogenates were centrifuged at 600 x g for 10 min 4°C. The remaining pellet was resuspended with the remaining supernatant and two additional lysing series were performed, with an in-between centrifugation step (600 x g for 10 min 4°C). After the first and last centrifugation, the remaining supernatant was recovered (corresponds to F1). The remaining pellet was resuspended in 800 μl MB and was centrifuged at 700 x g for 10 min at 4°C to obtain the crude mitochondria pellet and the second supernatant fraction (F2). Crude mitochondria pellets were resuspended in 250 μl MB and layered into a discontinuous sucrose gradient formed by a 6 ml layer of 1.2 M sucrose over a 5 ml layer of 1.6 M sucrose using ultra-clear tubes (14 x 89mm, Beckmann). Loaded gradients were centrifuged using a Beckman ultracentrifuged equipped with a SW40Ti rotor at 96000 x g for 2 hours at 4°C (acceleration and deceleration parameters were set to 3). Upon centrifugation, bands of interest were recovered. On the lighter sucrose layer, the band corresponding to membrane fraction (MMF) was collected and at the layer interface, the band corresponding to enriched mitochondria fraction (MTF) was collected. Upon collection, sucrose was diluted with four volumes of MB and all fractions were pelleted at 13000 x g for 10 min at 4°C. Pellets were resuspended in urea buffer (10 mM Tris, pH 8.0, 100 mM NaH2PO4, and 8 M urea). Equal protein amounts of each fraction were subjected to western blot analysis.

### Native Ni-NTA pulldown

Cells were pelleted at 9727 x g for 1 min at 4°C. Pellets were lysed with native lysis buffer (50 mM NaH_2_PO_4_, 300 mM NaCl, 0.2% NP-40, pH 8.0) complemented with 25 mM Imidazole, protease (Roche, 04693116001) and phosphatase inhibitors (Roche, 4906845001), respectively. Lysed pellets were incubated on ice for 30 min. After incubation, lysed pellets were centrifuged at 12000 x g for 15 min at 4°C and supernatant was collected for total protein quantification. Ni-NTA spin columns (Qiagen, 31014) were used to purify 6HIS-tagged TAX1BP1 and its interacting proteins following provider instructions. Briefly, Ni-NTA spin columns were first equilibrated using native lysis buffer with 25 mM Imidazole and centrifuged for 2 min at 890 x g. After equilibration, 250 µg of total protein were loaded on the columns and lysates were centrifuged at 270 x g for 10 min. Ni-NTA columns were washed three times with lysis buffer containing 50 mM imidazole, centrifuged for 2 min at 890 x g. After washing, pulldown fractions were collected with elution buffer (50 mM NaH_2_PO_4_, 300 mM NaCl, 0.2% NP-40, 500 mM Imidazole, pH 8.0) at 890 x g for 2 min twice. Samples were sent directly in elution buffer for mass spectrometry analysis.

### Ubiquitin pulldown

Cells were regularly washed with PBS before harvesting. After harvesting, cell pellets were centrifuged at 9727 x g for 1 min at 4°C. Pellets were resuspended in RIPA buffer (50 mM Tris/HCl, 150 mM NaCl, 1% NP-40, 0,5% deoxycholate, 0,2% SDS at pH 7.5) containing phosphatase and protease inhibitors and left on ice. After 30 min, homogenates were centrifuged at 18.000 x g for 15 min at 4°C and supernatant was recovered. Protein concentration was measurement using the Pierce BCA assay kit (ThermoFischer). Ubiquitin affinity beads (#UBA01B, Cytoskeleton) were used to pulldown ubiquitinated proteins. First, both ubiquitin affinity beads and control beads (#CUB02B, Cytoskelton) were reconstituted following manufacturer’s instructions. For each pulldown reaction, 20 µl of ubiquitin or control beads were aliquoted in a tube on ice and further washed twice with TBS-T. Following bead wash, 200 µg of total protein were loaded on the beads. Pulldown reaction tubes were filled with dilution buffer (50 mM Tris/HCl, 150 mM NaCl) containing protease and phosphatase inhibitors to reach 1.5 ml of volume and were left in a rotating platform for 2 h at 4°C. After incubation, beads were collected and washed with dilution buffer. Three washing steps were performed as follows: after buffer addition, tubes were allowed to rotate for 5 min at 4°C, further pelleted at 5.000 x g for 1 minute at 4°C and supernatant was removed. After the final wash, supernatant was completely removed with a Hamilton needle and 50 µl of reducing buffer was added instead. Samples were boiled for 10 min at 95°C. Bead pellets were further centrifuged at 5.000 x g for 1 minute at 4°C and supernatant was collected. Samples were further analyzed via Western Blot.

### Mass spectrometry

Eluates were mixed with 4x LDS sample buffer and proteins were separated by short SDS-PAGE runs using NuPAGE NovexR Bis-Tris 4–12% Mini Gels (Invitrogen, NP0335BOX). Protein bands were stained using QC Colloidal Coomassie Stain (Bio Rad Laboratories, 1610803) and whole sample lanes were excised. Gel pieces were further cut into small pieces prior to in-gel digestion. Gel pieces were destained by incubation with 5 mM ABC in 50% (v/v) ACN for 20 min three times and dehydrated using acetonitrile. Cysteine bonds were reduced by incubation with 10 mM DTT for 45 min at 65°C before carbamidomethylation by incubation with 55 mM iodoacetamide for 45 min at RT in the dark. Gel pieces were washed two times and dehydrated as before, followed by digestion with trypsin overnight at 37°C. Peptides were extracted in three consecutive incubation steps with Extraction buffer A (50% (v/v) ACN, 3% trifluoroacetic acid [TFA]), Extraction buffer B (80% (v/v) ACN, 0.5% (v/v) TFA) and acetonitrile for 30 min each. Supernatants of each extraction were pooled, and acetonitrile was evaporated by vacuum centrifugation. Peptides were purified using StageTips [58]. Mass spectrometric analysis was performed using an Orbitrap Exploris 480 online (Thermo Fisher Scientific, Germany) coupled to an EASY nLC-1200 system (Thermo Fisher Scientific, Germany). Peptides were separated on a 20 cm HPLC column with 75 µm inner diameter (CoAnn Technologies, ICT36007508F-50) in-house packed with 1.9 µm ReproSil-Pur C18 -AQ silica beads (Dr. Maisch HPLC GmbH, r119.aq.0001) and eluted by solvent B in a 60 min linear gradient from 5–33% at a flow rate of 200 nL/min. Carry-over was minimized by 30 min wash runs after each sample. Ionization was performed by ESI and the mass spectrometer was operated in positive ion mode controlled by XCalibur (Thermo Fisher Scientific). Acquisition of spectra was performed with a scan range of 350 – 950 m/z and a resolution on the MS1 level of 60 000. MS2 scans were performed using DIA isolation windows of 8 m/z with an overlap of 1 m/z (75 isolation windows), HCD collision energy of 30% and resolution of 15 000. The mass spectrometry proteomics data have been deposited to the ProteomeXchange Consortium via the PRIDE partner repository [59] with the dataset identifier PXD063982.

### Mass spectrometry analysis

Raw files were processed using Spectronaut version 19 in directDIA+ mode using optimized identification settings as described before [60]. Spectra were predicted using a Uniprot *Homo sapiens* database (104 556 entries, downloaded on 2024/01/30). The data were exported using the protein pivot report using PG.ProteinGroups as a unique protein ID and the PG.Quantity as a quantification column. Data analysis was performed using Perseus version 2.0.5.0 [61], Microsoft Excel and R. Proteins were annotated functionally and by compartment using Gene Ontology Biological Processes and Gene Ontology Cellular Compartment databases (both downloaded on 2024/10/29) as well as for mitochondrial sub-compartments using MitoCarta3.0 (downloaded on 2021/02/24). To assess differences between WT and mutant TAX1BP1, student’s T-tests were performed on protein intensities normalized to the protein quantities of the empty vector for both CCCP only and CCCP and BafA treated samples. Significantly up- and downregulated proteins (p<0.05) were further analyzed for statistical overrepresentation on GO biological processes using the panther database (pantherdb.org, version 19.0) [23].

### CellProfiler-based image analysis

The version 4.2.1 of the CellProfiler software was used [62]. All analyses rely on pooled single cell values obtained from at least three independent biological replicates. In general, individual cells were initially recognized with the nuclear staining and a cytoplasm area of 80-200 pixels (px) around the nucleus was further categorized as the cytoplasm, unless otherwise noted. Areas of interest per cell correspond to the established cytoplasm without the nucleus area.

For the analysis of mitochondrial clustering in WT-Parkin HeLa transfected with His-empty or HIS-tagged TAX1BP1, mitochondrial material was recognized using SSBP1 signal as primary object. SSBP1 objects were identified using a robust background two-class thresholding method with lower and upper threshold bounds of 0.0-1.0, an adaptive window of 50 and a size range of 6-500 px. Next, single SSBP1 objects that were in close contact were merged (distance between objects=0 px). To account for only positively TAX1BP1 transfected cells, TAX1BP1 was recognized as primary objects using a minimum cross entropy two-class thresholding method with a lower and upper threshold bounds of 0.0-1.0, an adaptive window of size 30 and a size range of 10-40 px. Only cells with positive TAX1BP1 signal (count TAX1BP1 objects>0) were taken into account for the TAX1BP1 overexpressing (OE) condition.

For the analysis of mitochondria clustering in 5KO HeLa co-expressing parkin as well as WT-, K549R and TAX1BP1ΔCC3, the analysis was performed similarly as previously described for the WT-Parkin HeLa with some variations. Mitochondrial material was recognized as SSBP1 signal using a roust background two-class thresholding method with lower and upper threshold bounds of 0.002-1.0, an adaptive window of size 50 and a size range of 4-500 px. TAX1BP1 was recognized as primary objects using a minimum cross-entropy two-class thresholding method with lower and upper threshold bounds of 0.0022-1.0, an adaptive window of size 80 and a size range of 6-40 px. Similarly, Parkin expression was monitored by recognizing parkin signal as primary object using a minimum cross-entropy two-class thresholding method with lower and upper threshold bounds of 0.0015-1.0, an adaptive window of size 50 and a size range of 10-40 px. Parkin objects were further merged if they were in close contact (object distance=0 px). Only cells with positive TAX1BP1 and parkin signal (mean object area > 0) were taken into account for this analysis. Values of total mean SSBP1 area and mitochondrial area or perimeter normalized by cell size (area or perimeter) are shown. For the analysis of TAX1BP1 distribution on SSBP1 clusters, only cells with a mean TAX1BP1 intensity of ≥10 aiu (arbitrary intensity units) were taken into account and the recognition method of TAX1BP1 and parkin signals was optimized. In this case, TAX1BP1 and parkin were recognized as primary objects using a robust background two-class thresholding method with lower and upper threshold bounds of 0.0035-1.0 (for TAX1BP1) or 0.002-1.0 (for parkin) with a size range recognition of 3-40 px in both cases. Mitochondria were recognized using SSBP1 signal as described before for 5KO HeLa. To measure the overlapping area between TAX1BP1-SSBP1 and parkin-SSBP1, total SSBP1 area was masked with TAX1BP1 or parkin merged objects and the percentage of SSBP1 overlapped by each protein was calculated (% SSBP1 recognized). Similarly, percentages of TAX1BP1 or parkin on mitochondria were calculated by considering which portion of the total TAX1BP1 or parkin area was engaged with mitochondrial material (% TAX1BP1 or parkin on SSBP1).

For the analysis of size classification of TAX1BP1^+^ puncta and TAX1BP1-HSP60 material overlap in WT-Parkin (a) and 5KO-PRKN HeLa (b), cell area was recognized using a watershed-image method with nuclei as a reference point and TAX1BP1 or Parkin signal to identify cell contours (minimum cross entropy method, threshold bounds: 0.00001-1). TAX1BP1 puncta were recognized as primary objects using robust background method with lower and upper threshold bounds of 0.000001-1.0 in (a) and 0.0015-1.0 in (b), an adaptive window of size 15 and a size range of 4-40 px. In WT-Parkin Hela, only cells with TAX1BP1 mean intensity of 0.004 aiu were considered. To optimize the recognition of individual puncta, clumped objects were distinguished by shape but dividing lines between clumped objects were intensity-based. TAX1BP1 objects were merged when in contact with each other (distance = 0 px), this step allowed to merge TAX1BP1 puncta that were in close contact with other TAX1BP1 puncta and were particularly difficult to recognize as single objects. Upon merging TAX1BP1 single objects, objects with an area form factor of 0.68-

0.8 were kept (form factor=1 for a perfectly circular object). This step allowed to keep relatively circular-shaped objects (individual puncta) and discard merged objects that were the result of multiple TAX1BP1 puncta. Thus, not all TAX1BP1 puncta were taken into account for this analysis. After filtering, TAX1BP1 puncta were classified based on their size: enlarged puncta (area>150 px^2^), intermediate puncta (area≤150 px^2^) and small puncta (area<50 px^2^). Mitochondrial material (HSP60) was recognized as primary object using a robust background method with lower and upper threshold bounds of 0.001-1 and a size range of 1-40 px in (a) and 0.0018-1.0 and a size range of 4-200 px in (b) and a size. HSP60 objects were used to mask enlarged, intermediate and small TAX1BP1 puncta. In WT-Parkin Hela, all TAX1BP1 puncta sizes were classified between HSP60-positive or HSP60-negative according to their overlap. Percentages of HSP60 puncta co-localizing with overall or enlarged TAX1BP1 puncta are shown as well as the corresponding HSP60 puncta area co-localizing with enlarged TAX1BP1 puncta. For each analyzed image of this analysis, output images were generated, showing the localization of the classified TAX1BP1 puncta, the cell limits. Colocalization analysis in 5KO-PRKN was performed using the MeasureColocalization module in CellProfiler, where Pearson’s correlation coefficient of TAX1BP1 and HSP60 signals were computed per cell.

Colocalization analysis of TAX1BP1-HSP60 and LC3B-HSP60 in 5KO-PRKN HeLa was performed similarly as described before with some variations. To ensure the analysis only included cells with similar TAX1BP1 expression, cells with a minimum mean integrated intensity of 7 aiu were considered. A TAX1BP1 background region was established to filter out cells with excessive TAX1BP1 levels (max. integrated intensity of 20 aiu overlapping nucleus). HSP60 objects were recognized as previously described. TAX1BP1 objects were recognized using robust background method with lower and upper threshold bounds of 0.0022-1.0 and a size of 1-100 px. LC3B was recognized using a three-class Otsu method. Co-localization parameters were measured per each recognized cells for all three channels and Pearson’s coefficient as well as thresholded Mander’s coefficient were obtained.

### Statistics

All data is derived from at least three independent biological replicates unless otherwise indicated. Statistical analysis and graph depiction of the data was performed using GraphPad Prism 8.0.2. Statistical significance was established at p<0.05 unless otherwise indicated. Significance was assessed by using a one-way ANOVA for multiple comparisons and subsequent Tukey’s post-hoc test unless otherwise indicated. For the correlation analysis of TAX1BP1 binding to depolarized mitochondria, a Pearson correlation test was performed. For analysis in which only two conditions were compared, significance was assessed with an unpaired t-test analysis. The following significance indicators were used: ****: p<0.001, ***: p<0.005, **: p<0.005, *: p<0.05

## Acknowledgements

PentaKO (5KO) HeLa cells were kindly provided by Dr. Chunxin (Black) Wang and Dr. Richard J. Youle, NINDS. This work was supported by the DFG Research Training Group 2364 “MOMbrane”, the Hertie Foundation and the German Center for Neurodegenerative Diseases. We would also like to acknowledge Prof. Dr. Tassula Proikas-Cezanne for valuable discussion and feedback. Lastly, we would also like to acknowledge Henrietta Lacks and her family.

## Data availability

All mass spectrometry proteomics data have been deposited to the ProteomeXchange Consortium via the PRIDE partner repository and are openly available in http://www.ebi.ac.uk/pride with the dataset identifier PXD063982.

## Abbreviations

5KO: penta knock-out HeLa
ATG9A: autophagy-related protein 9A
BafA: Bafilomycin A
CALCOCO2/NDP52: calcium binding and coiled-coil domain 2
CC3: third coiled-coil domain of TAX1BP1
CCCP: carbonyl cyanide m-chlorophenylhydrazone
CD63: CD63 antigen
COX4: cytrochrome c oxidase subunit 4
CS: citrate synthase
HSP60: heat shock protein 60
LC3B: microtubule-associated protein 1 light chain 3 beta
MFN1: mitofusin-1
MMF: membrane fraction
MTF: mitochondria fraction
MYO6: unconventional myosin-VI
NBR1: NBR1 autophagy cargo receptor
OMM: outer mitochondrial membrane
OPTN: optineurin
SQSTM1/p62: sequestosome-1
PD: Parkinson disease
PINK1: serine/threonine-protein kinase PINK1
PRKN: E3-ligase parkin
PTM: post-translational modification
RAB7: ras-related protein rab-7a
SSBP1: single-stranded DNA-binding protein
TAX1BP1: Tax1-binding protein 1 homolog
TBC1D15: TBC1 domain family member 15
TOM20: translocase of outer membrane 20 kDa subunit
UBD: ubiquitin binding domain
VDAC: voltage-dependent anion channel
VPS35: Vacuolar protein sorting-associated protein 35
WDR45B: WD repeat domain phosphoinositide-interacting protein 3
WT: wild-type.

