## Supplementary figures for "Parkin-dependent ubiquitination of TAX1BP1 affects the degradation pathway of defective mitochondria"

### Figure S1

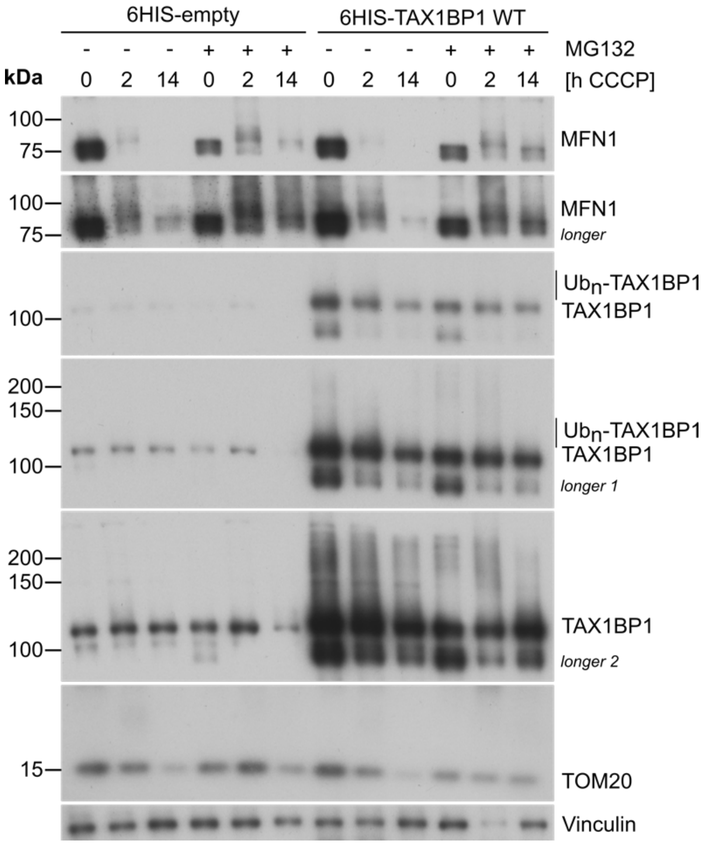


**Figure S1:** TAX1BP1 ubiquitination does not direct the autophagy adaptor to proteasomal degradation. WT-Parkin HeLa cells were transfected with HIS-tagged TAX1BP1 and subjected to mitochondrial depolarization for the indicated time points, in the presence (+) or absence (-) of the proteasomal inhibitor MG132. Mitophagy was confirmed by probing for the immediate-early proteasome substrate MFN1 and the later autophagy substrate TOM20. Vinculin served as loading control.

### Figure S2

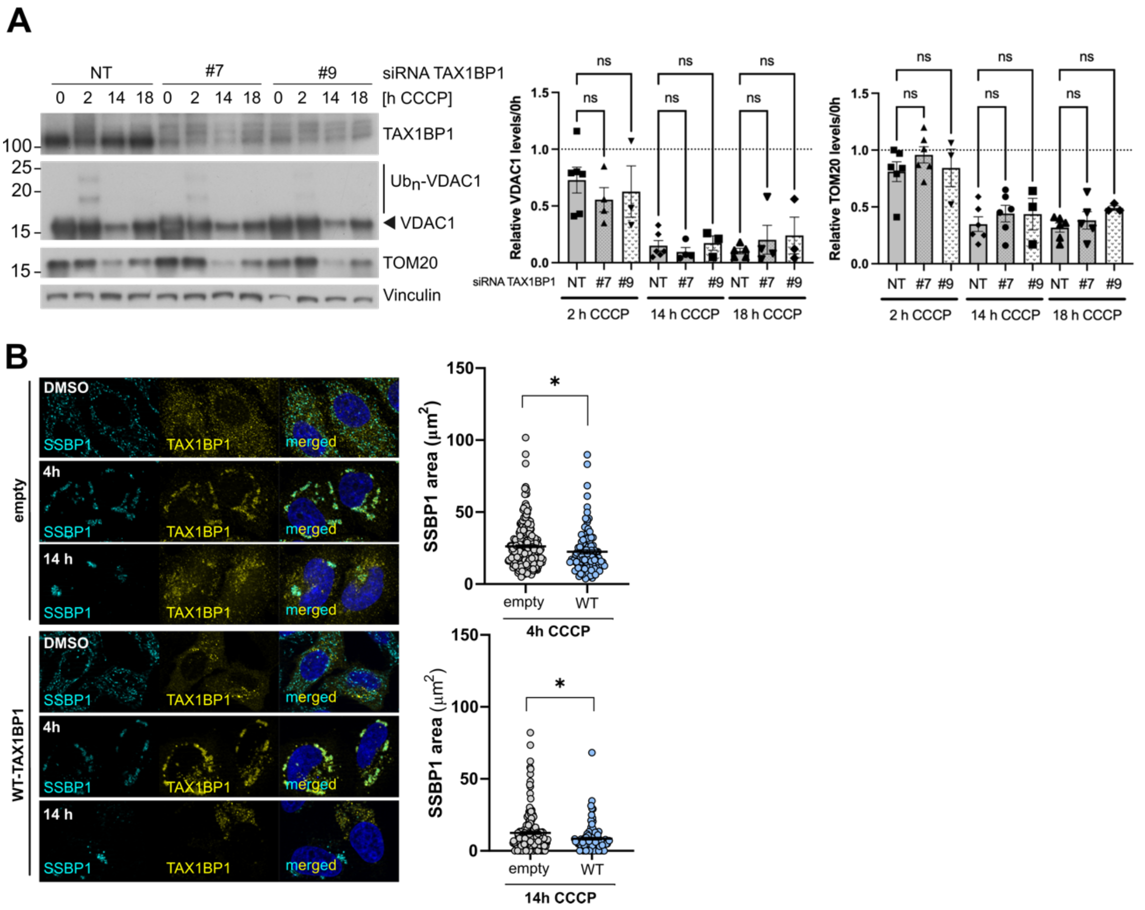


**Figure S2:** (**A**) TAX1BP1 siRNA mediated knockdown in WT-Parkin HeLa cells does not impair VDAC and TOM20 degradation upon acute mitochondria depolarization. WT-Parkin HeLa cells were transfected with siRNA against TAX1BP1 for three consecutive days. After the third day, mitochondria were depolarized with CCCP for the indicated time points and protein quantification was performed. (**B**) WT-Parkin HeLa cells were transfected with His-tagged TAX1BP1 for 24 h. Further, mitochondria were depolarized with 10 µm CCCP for the indicated time points. Left panel: Representative immunofluorescence images showing SSBP1 (cyan) and TAX1BP1 (yellow) upon mock or HIS-TAX1BP1 transfection after CCCP-mediated depolarization at the indicated time points. Right panel: Reduction of SSBP1 area after TAX1BP1 transfection after short (4 h) or extended (14 h) CCCP treatment. Scale bars: 5 µm.

### Figure S3

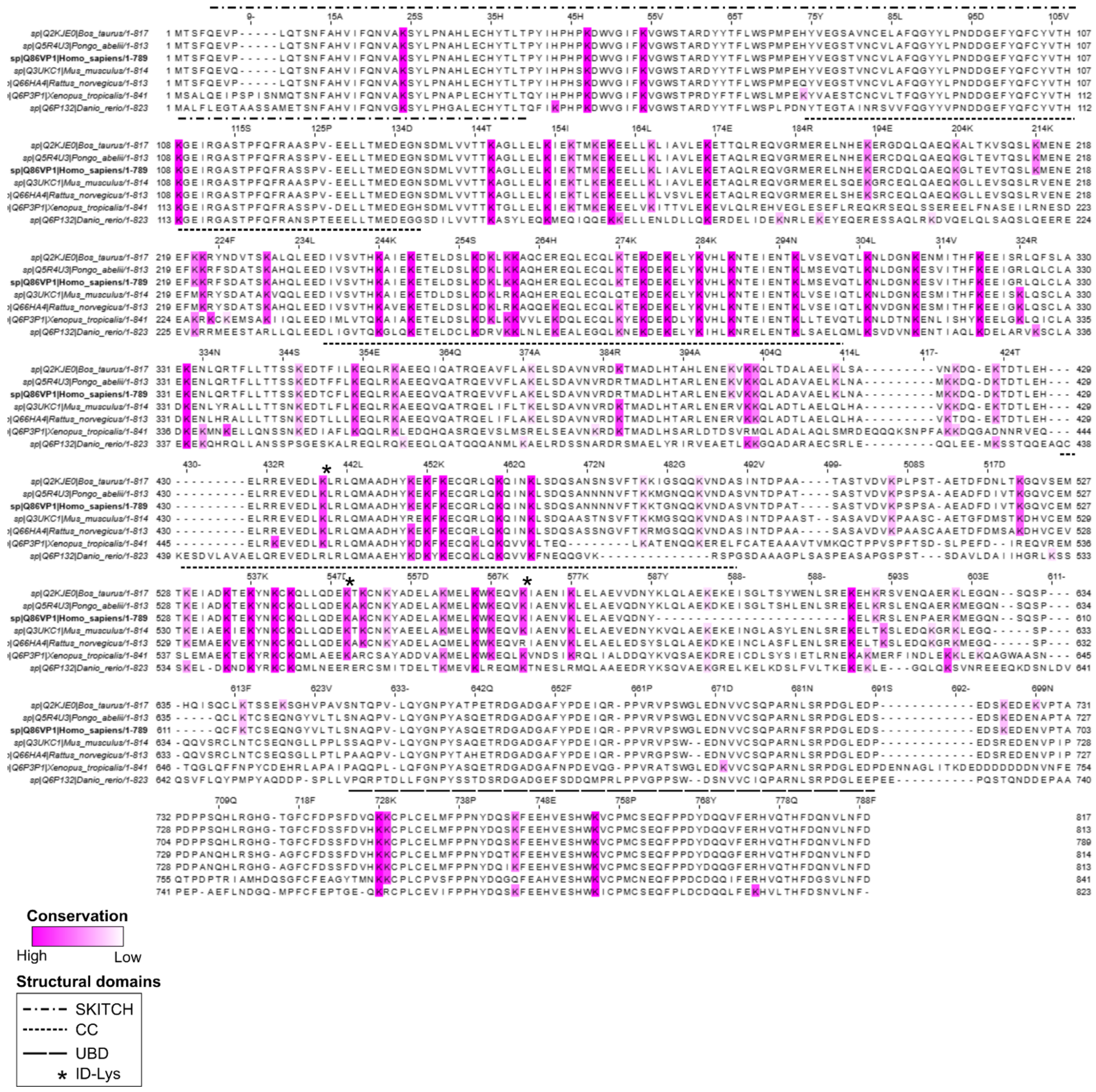

**Figure S3:** TAX1BP1 alignment across species with emphasis on lysine conservation. TAX1BP1 sequences from human (*Homo sapiens*), cow (*Bos Taurus*), orangutan (*Pongo abelii*), mice (*Mus musculus*), rat (*Rattus norvegicus*), frog (*Xenopus tropicalis*) and zebrafish (*Danio rerio*) were aligned using the online tool MAFFT (https://mafft.cbrc.jp/alignment/server/index.html) and the software JalviewJS (Version 2.11.4.1) was used for visualization and lysine conservation analysis [63,64]. All lysine residues are shown in magenta across all TAX1BP1 species. (*) Identified lysines in the mass-spectrometry screen. Structural domains are represented with the indicated symbol lines. Lysine conservation analysis was performed with JalviewJS software, using the conservation color increment function with a threshold of 25.

### Figure S4

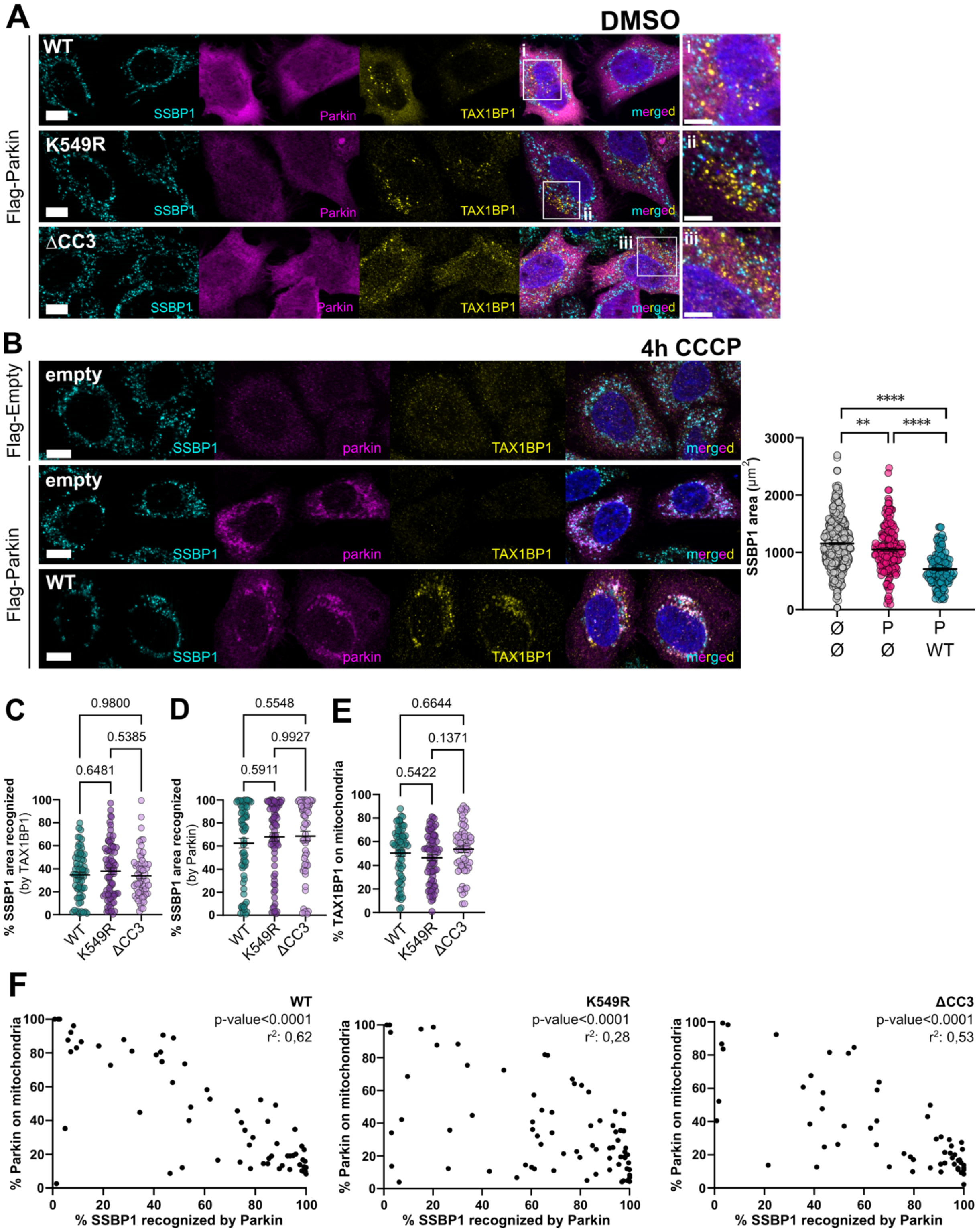


**Figure S4:** (**A**) Overexpression of HIS-tagged WT, TAX1BP1$^{K549R}$ or TAX1BP1ΔCC3 (yellow) in 5KO-HeLa cells co-transfected with Flag-parkin (magenta). Mitochondria are stained with SSBP1 (cyan). Scale bars: 10 µm. (**B**) Left panel shows representative images of 5KO-HeLa cells stained for SSBP1 (cyan), overexpressing Flag-parkin (magenta) or Flagempty control plasmid in combination with His-tagged WT-TAX1BP1 (WT) (yellow) or its respective empty control plasmid (empty). Cells were treated with 10 μM CCCP for 4 h. Right panel shows CellProfiler based quantification of mitochondrial area represented by SSBP1. Ø: empty plasmid. P: 3xFlag-Parkin co-transfection. Scale bars: 10 μm. (**C-F**) CellProfiler based quantification of several parameters of 5KO HeLa cells co-transfected with Flag-parkin and TAX1BP1 (WT and mutants) after 4h CCCP. (**C-D**) Percentage of SSBP1 area recognized by (**C**) TAX1BP1 and (**D**) Parkin. (**E**) Percentage of TAX1BP1 found on mitochondria after 4h CCCP. (**F**) Pearson correlation analysis of the percentage of translocated parkin (Y) and the percentage of SSBP1 area recognized (X) in cells co-transfected with WT, TAX1BP1$^{K549R}$ and TAX1BP1$^{\Delta CC3}$.

### Figure S5

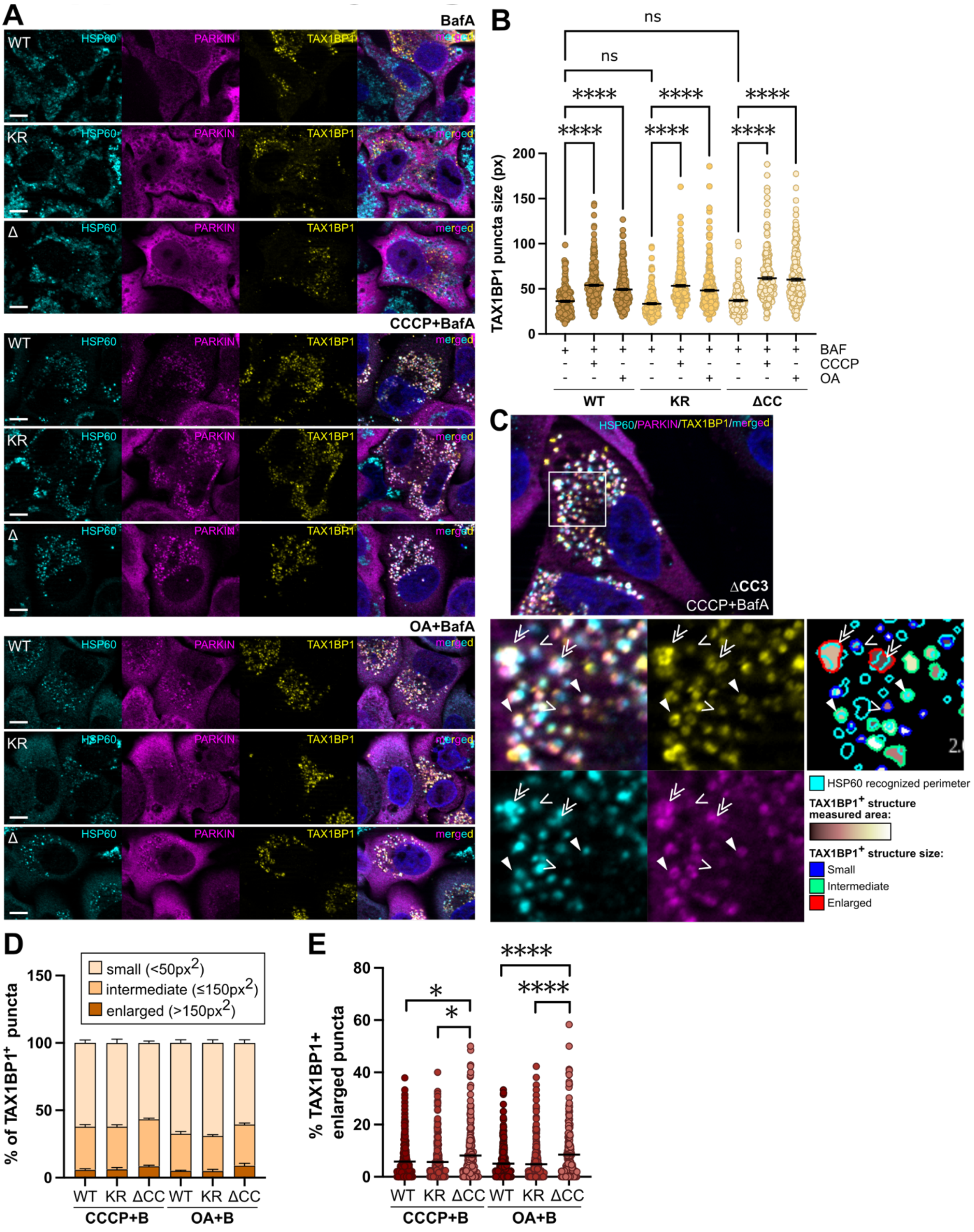


**Figure S5:** (**A**) Representative images showing TAX1BP1 (yellow), Parkin (magenta) and HSP60 (cyan) in 5KO-PRKN HeLa transiently expressing WT, TAX1BP1$^{K549R}$ (KR) or TAX1BP1ΔCC3 (ΔCC) and further treated for 16 h with BafA alone or in combination with CCCP or OA. (**B**) TAX1BP1 puncta size under the indicated treatment and transfection conditions. (**C**) Example image of the

CellProfiler-based identification of enlarged (double arrowheads or red), intermediate (thick arrowheads or green) or small (thin arrowheads or blue) TAX1BP1 structures in 5KO-PRKN HeLa cells overexpressing TAX1BP1ΔCC3 after CCCP combined with BafA treatment. Structures with colored areas indicate identified TAX1BP1 puncta, the area of which was measured and further used for classification. (**D**) Classification of TAX1BP1$^+$ structures observed after the indicated treatment and transfection conditions. (**E**) Percentage of enlarged TAX1BP1 puncta identified.

### Figure S6

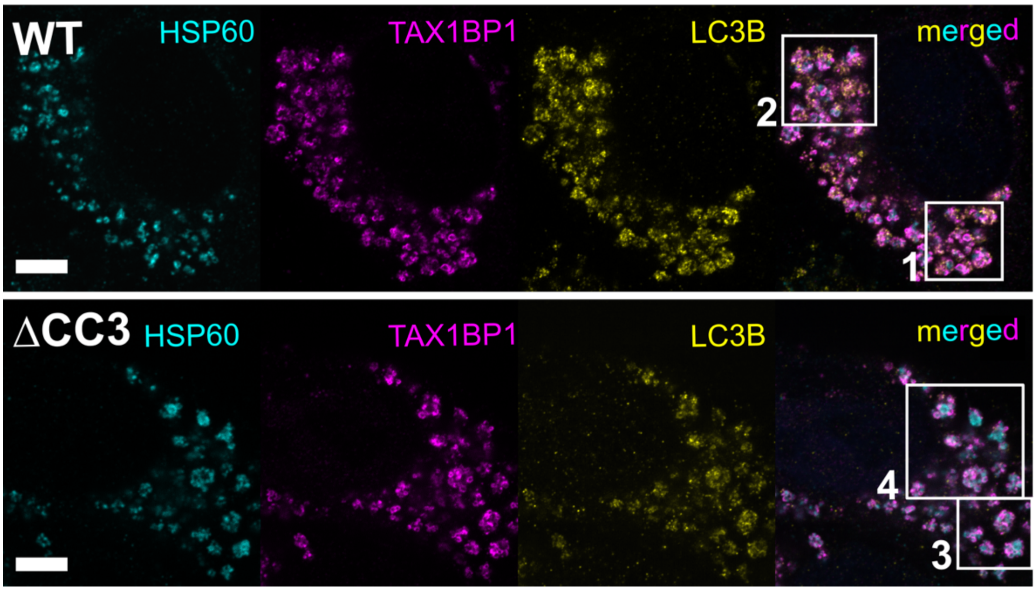


**Figure S6:** Enlarged vesicles found in WT and delCC3 upon CCCP and BafA treatment (16 h). Representative pictures shown in main Figure 6A where two different regions are selected for each condition (WT and TAX1BP1ΔCC3). A full z-stack is provided for the selected regions in video format (see Supplementary videos 1-4, respectively).

### Figure S7

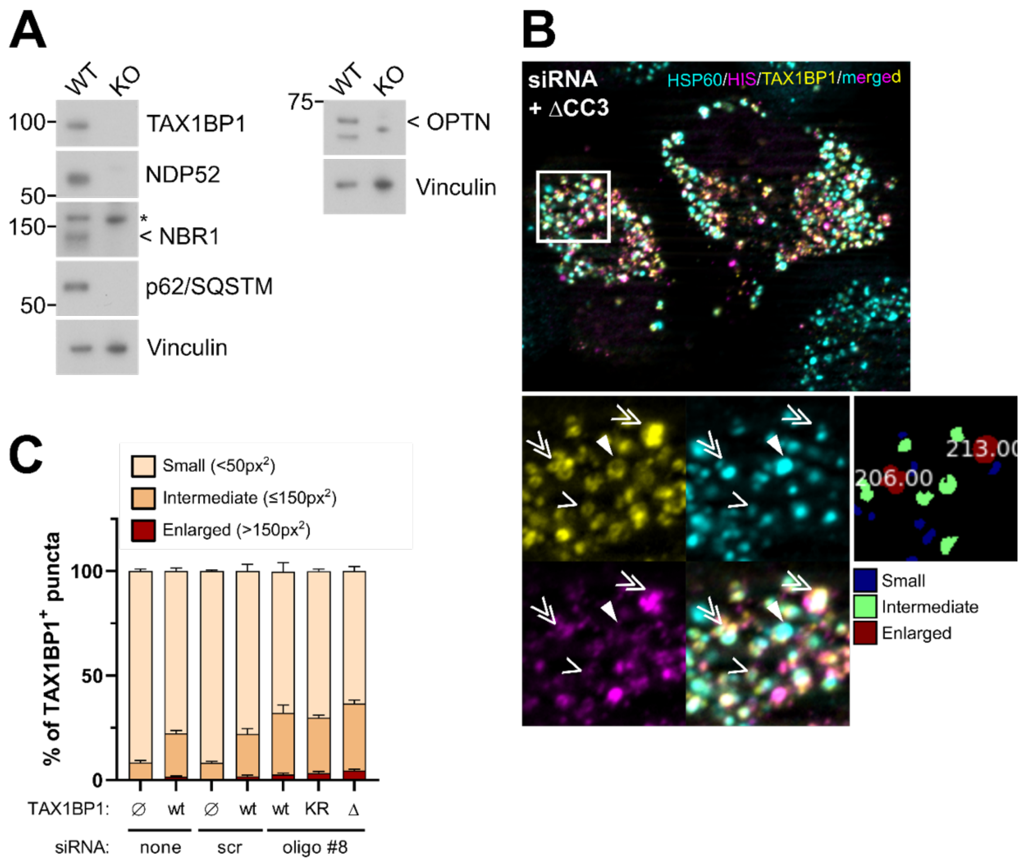


**Figure S7:** Characterization of TAX1BP1$^{+}$ vesicles in WT-Parkin HeLa cells after transient knockdown and re-transfection. (**A**) Protein expression of classical mitophagy associated autophagy adaptors in WT-Parkin and 5KO-PRKN HeLa cells. Representative images of two independent biological replicates (n=2). (**B**) Example image of the CellProfiler-based identification of enlarged (double arrowheads), intermediate (thick arrowheads) or small (thin arrowheads) TAX1BP1 structures in WT-Parkin HeLa cells overexpressing TAX1BP1ΔCC3 after 16h CCCP combined with BafA. Numbers in white indicate pixel area of TAX1BP1$^{+}$ enlarged structures. (**C**) Classification of TAX1BP1$^{+}$ puncta observed in WT-Parkin HeLa cells after siRNA mediated knockdown, CCCP-mediated depolarization combined with BafA treatment.

### Figure S8

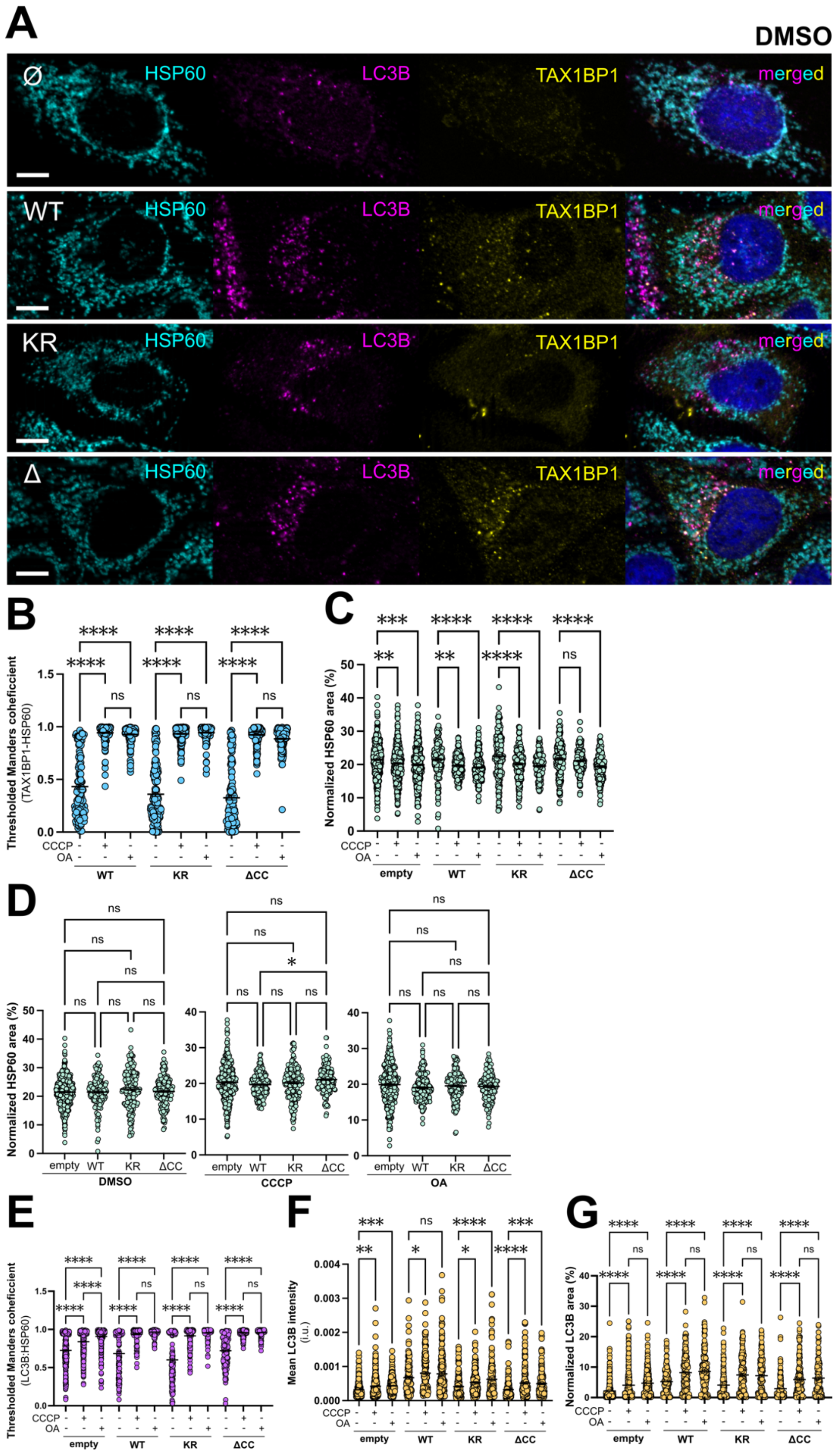


**Figure S8:** Co-localization analysis of TAX1BP1 and LC3B on mitochondria material under control conditions (DMSO) or after 4 h depolarization with CCCP or OA in 5KO-PRKN HeLa cells transiently expressing HIS-tagged TAX1BP1 WT, K549R (KR) or ΔCC3 (ΔCC). (**A**) Representative images showing TAX1BP1 (yellow), LC3B (magenta) and HSP60 (cyan) under control conditions. (B) Thresholded manders coheficcient of TAX1BP1-HSP60 signal. Scale bars: 8 µm. (**C**-**D**) Changes in HSP60 area upon CCCP- or OA-mediated depolarization for each TAX1BP1 overexpressing condition. Treatment comparisons are shown in panel (C) while TAX1BP1 transfection effect is shown in panel (D) for each individual treatment condition. (**E**) Thresholded manders coheficcient of LC3B-HSP60 signals. (**G**) Mean LC3B intensities across all treatment and TAX1BP1 transfection conditions. (**H**) Normalized LC3B area relative to whole cell area. In all panels, data of individual cells is represented by single dots. All data come from at least three independent experiments.

### Figure S9

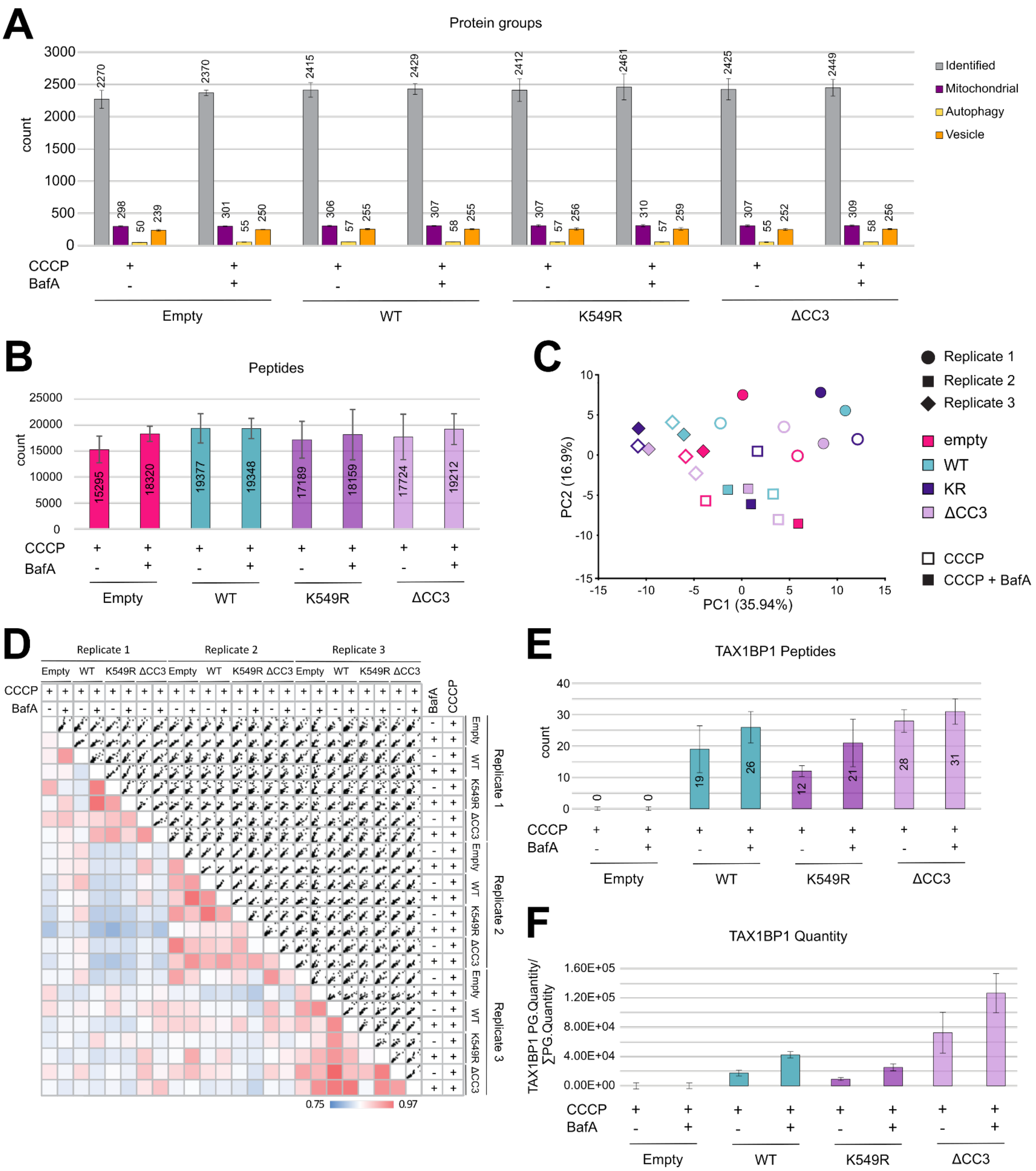


**Figure S9:** Quality control of the native pulldown mass spectrometry analysis. (**A**) Median of the number of protein groups identified per sample alongside their portion affiliated to mitochondria, the autophagy or the vesicular transport machineries. (**B**) Median of the number of peptides identified per sample. (**C**) Principal component analysis of all samples with each replicate treated individually. (**D**) Spearman rank correlation of PG Quantity with each replicate treated individually. (**E**) Median of the number of TAX1BP1 peptides per sample. (**F**) Median of the protein abundance of TAX1BP1 normalized to the total protein intensity per sample.

### Figure S10

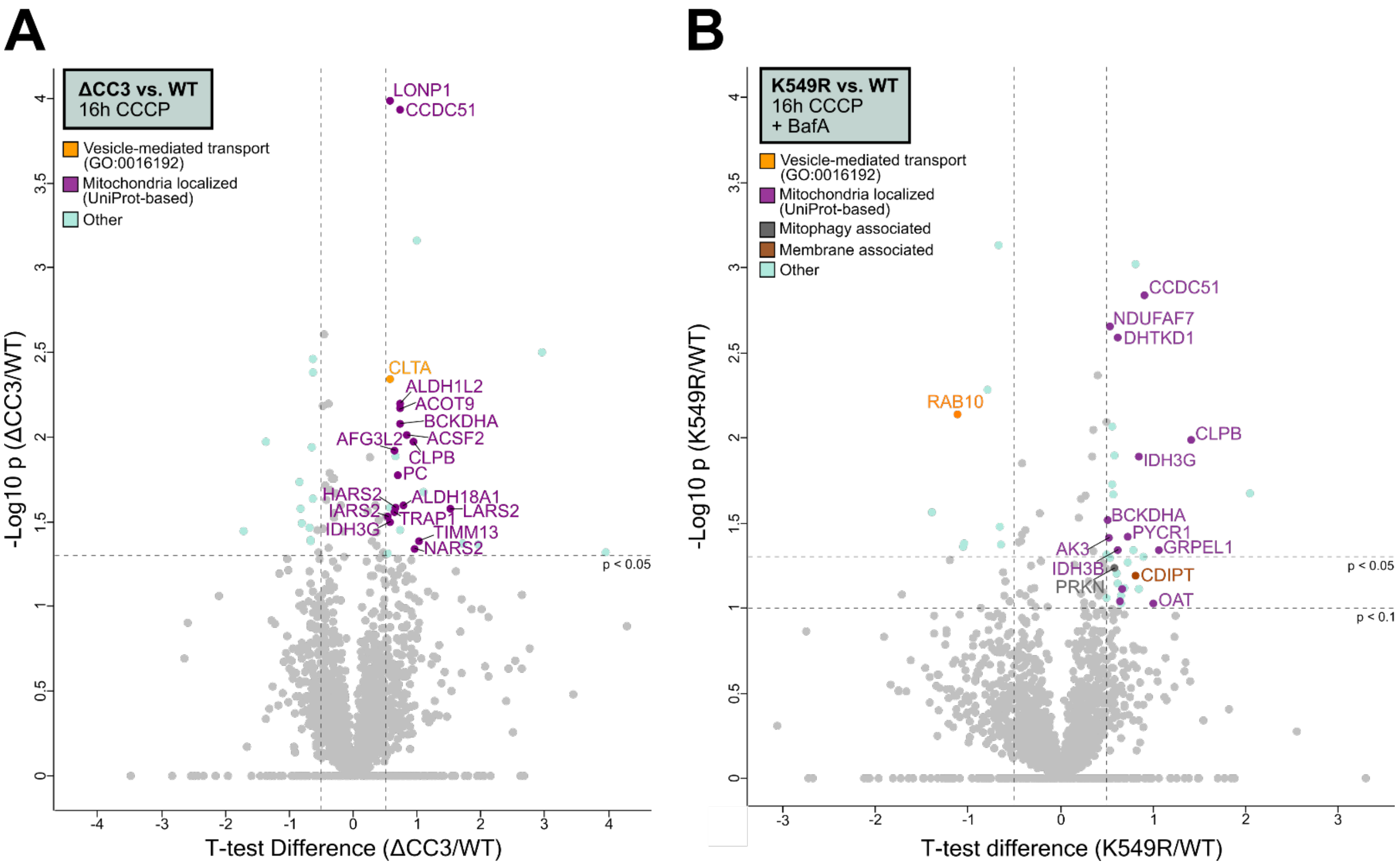


**Figure S10:** Volcano plots showing TAX1BP1$^{K549R}$ and TAX1BP1ΔCC3 after CCCP treatment only or treatment combined with BafA. (**A**) Statistical analysis of TAX1BP1ΔCC3 vs. WT TAX1BP1 after extended mitochondrial depolarization. (**B**) Statistical analysis of TAX1BP1$^{K549R}$ vs. WT after CCCP treatment combined with BafA. In both cases, data are represented by the $\log_{10}$ p-value plotted against student's T-test difference. Significance lines represent p-value $\leq 0.05$ and p-value $\leq 0.1$, as indicated. Mitochondria localized proteins are highlighted in lilac and were annotated according to UniProt. Other annotations correspond to Vesicle-mediated transport (GO:0016192) in orange, proteins related to mitophagy (gray) or proteins associated with membranes (brown).
